# Summary Statistics and Approximate Bayesian Computation Are Comparable to a Common Convolutional Neural Network Design for Inferring Fixation Times

**DOI:** 10.64898/2026.02.17.706432

**Authors:** Miles D. Roberts, Emily B. Josephs

**Affiliations:** Genetics and Genome Sciences Program, Michigan State University, East Lansing MI; Department of Plant Biology, Michigan State University, East Lansing, MI; Ecology, Evolution, and Behavior Program, Michigan State University, East Lansing, MI; Plant Resilience Institute, Michigan State University, East Lansing, MI

**Keywords:** Machine Learning, Approximate Bayesian Computation, Selective sweeps, Time to fixation

## Abstract

Detecting signatures of positive selection in genomes is a common application of population genetics and one of the most influential models for this task is the hard selective sweep where a *de novo* mutation rapidly fixes. Many statistics aim to detect hard sweeps by summarizing signatures left behind in the site frequency spectrum, linkage disequilibrium, and haplotype frequencies. However, potentially undiscovered signals could still exist. We attempted to test whether any undiscovered signatures of hard sweeps exist by comparing a convolutional neural network, which can learn signatures from raw data without any prior knowledge, to known summary sweep statistics for inferring the time to fixation (*t_f_* ) of a hard sweep in a background of variable sweep ages (*t_a_*). Across approximately 250,000 simulations encompassing 5 different demographic scenarios of single panmictic populations, a convolutional neural network model trained directly on raw genotype data failed to better predict *t_f_* than methods based purely on common summary statistics. This suggests that few undiscovered signals remain in single timepoint, single population genotype data to better predict *t_f_* . However, future models can explore additional demographic spaces and penalize use of known population genetic features to potentially improve this search.

## Introduction

How long does it take for beneficial alleles to spread? This question was at the top of mind of naturalists in the early 20th century as there was doubt over natural selection’s importance as an evolutionary mechanism (see Charlesworth [2020] for a fuller historical treatment). The thinking was that if selection increases the frequency of beneficial alleles only very slowly, then maybe natural selection is not important in shaping divergence. However, early models showed that while most beneficial mutations are lost by drift [Fisher, 1923, Haldane, 1927] the length of evolutionary timescales and rate of new mutations are high enough that some beneficial mutations eventually fix [Zhao et al., 2013]. The time required (usually in generations) for a *de novo* mutation to fix is called the time to fixation (*t_f_* ). As it became appreciated that *t_f_*was short enough for natural selection to be a feasible evolutionary mechanism, further studies of *t_f_* were driven by interest in fixation of neutral mutations, including studies of molecular substitution rates [Kimura and Ohta, 1969] and trait degradation as a result of relaxed selection [Kimura, 1980]. The theories from this foundational work have since been extended to understand how *t_f_* is shaped by factors such as changing environments [Cui and Yuan, 2018, Kaushik and Jain, 2021], population structure [Greven et al., 2016].

Despite this foundation, studies of *t_f_* for beneficial mutations have subsided in recent years [Zhao et al., 2013, Charlesworth, 2020]. It is instead much more common for studies to estimate coalescent times [Brandt et al., 2022], variant age [Bisschop et al., 2021], or allele frequency trajectories of currently segregating alleles [Stern et al., 2019]. One reason for this is that *t_f_* is tightly correlated to the selection coefficient (*s*) in simple models. For example, the mean *t_f_* for a single additive beneficial mutation in a constant-sized population of *N_e_* individuals is 2 ln(2*cN_e_* 1)*/s* where *c* is ploidy level [Otto and Whitlock, 2013]. This equation is most sensitive to *s* because it is outside the logarithm. Thus, under this model one could study *s* and get essentially the same information as *t_f_* . However, the exact relationship between *t_f_* and *s* is more complicated when arbitrary dominance, mating system variation, and demographic changes are considered [Glémin, 2012]. Selection coefficients and coalescent times thus do not fully convey the timescale of fixation by themselves outside of simple scenarios. Given a putative sweep, estimates of *t_f_* help summarize the evolutionary timescale for a focal mutation or trait [Zhang et al., 2009].

How can we estimate *t_f_*from genetic data of a sweep site? Hard sweeps have long been known to leave behind clear genomic signatures [Smith and Haigh, 1974, Kaplan et al., 1989, Stephan et al., 1992, Stephan and Mitchell, 1992, Barton, 1998]. The work of Smith and Haigh [1974] outlined how sweeps result in a signature of reduced heterozygosity and a tract of complete homozygosity around the sweep site and that the width of this tract depends on the recombination rate and strength of selection. These “dips” in diversity can be used to fit simple analytical models as a function of *t_f_* , assuming a single *de novo* additive mutation that is sampled immediately after sweep completion [Coop and Ralph, 2012, Coop, 2020]. If the sweep age (*t_a_*, the time between reaching high frequency and sampling, Figure 1A) is small then any degradation of sweep patterns is due mainly to variation in *t_f_* . However, if *t_a_*is large then different combinations of *t_f_*and *t_a_* can create the same diversity patterns - a statistical situation called non-identifiability. For instance, sweeps that fixed recently (low *t_a_*) but took a long time to fix (high *t_f_* ), leave similar signatures as sweeps that fixed rapidly (small *t_f_* ) but longer ago (large *t_a_*, Figure 1C). It is possible to better disentangle *t_f_* for arbitrary *t_a_* by applying more complex analytical models [Schraiber et al., 2016, He et al., 2020, Bisschop et al., 2021], but these approaches will also reach non-identifiability eventually if *t_a_* is large enough. This poses the question of how much undiscovered signal remains in polymorphism data to infer sweep properties like *t_f_*[Bisschop et al., 2021].

**Figure 1.**
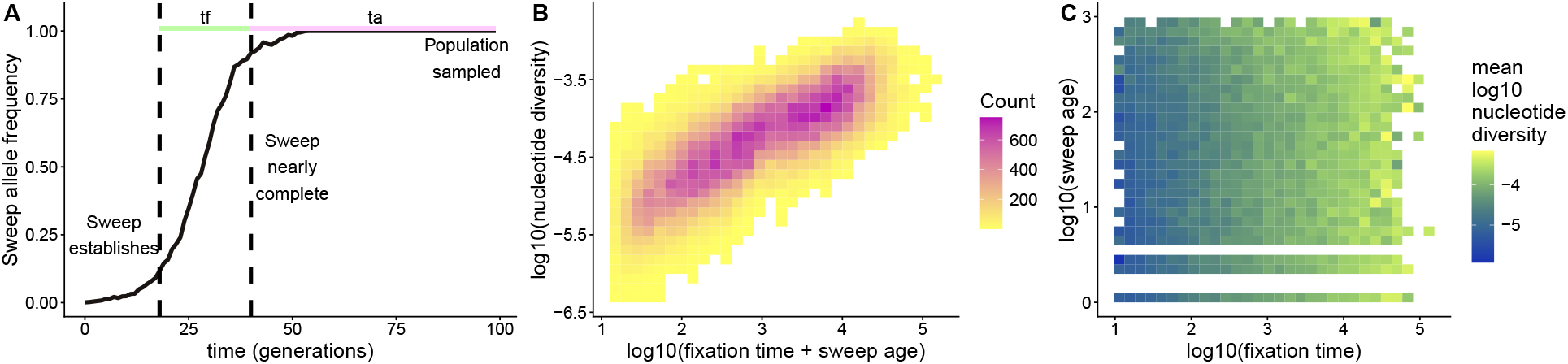
Separately estimating *t_f_* and *t_a_* from summary statistics leads to unidentifiability. (A) Example allele frequency trajectory comes from simulating a beneficial allele with selection coefficient 0.5, dominance 0.5, and initial frequency of 1/1000 in a Wright-Fisher population of size 1000. (B) Nucleotide diversity in the sweep region linearly scales with the total lifetime of the sweep: *t_f_* + *t_a_*. (C) The total *t_a_* is capped to be *<* 0.1*N_A_* based on the initial range of simulations from **Batch 1** (Figure S1). Thus young, slow sweeps (high *t_f_*and low *t_a_*) can be distinguished from old, fast sweeps (low *t_f_* and high *t_a_*).

Instead of estimating *t_f_* with analytical models, which typically involve making a fixed set of assumptions (Wright-Fisher populations, a range of *t_a_*, etc.), one could instead leverage evolutionary simulations [Haller and Messer, 2019, Haller et al., 2019, Baumdicker et al., 2022]. The core idea is to simulate evolutionary scenarios that reasonably reflect the study system, and then build a model that predicts an unobserved evolutionary parameter (e.g. *t_f_* ) from the simulated population’s genetic data [Bertorelle et al., 2010, Murga-Moreno et al., 2023]. Overall, this paradigm of training models on simulated data offers incredible flexibility and allows one to match the assumptions of a model to the current understanding of the study system. However, multiple frameworks for predicting unobserved parameters from simulated data are available. One common simulation-based framework in evolutionary genetics is Approximate Bayesian Computation (ABC), which builds regression-like models where the explanatory variables are pre-defined summary statistics (e.g. *π*, Tajima’s D, etc.) calculated from simulated populations and the outcome variable is the evolutionary parameter of interest [Beaumont et al., 2002, Csilĺery et al., 2010]. Once these relationships are deduced, the models can then be applied to summary statistics calculated from real data to infer an unobserved evolutionary parameter. Examples of parameters estimated with ABC include the timing of selection on beneficial mutations [Przeworski, 2003, Ormond et al., 2016, Nakagome et al., 2019], demographic rates [Pascual et al., 2007, Bertorelle et al., 2010], admixture proportions [Robinson et al., 2014], and the distribution of fitness effects [Johri et al., 2020, 2022]. Despite the broad applicability of ABC, the need to compute pre-defined summary statistics can be a draw-back because a researcher may not know every feature *a priori* that is important to their prediction problem.

A second method for building inferences on simulated data is machine learning (ML). In contrast to ABC, ML models learn important prediction features directly from data, making them potentially able to discover features that the researcher would not think to pre-calculate [Sheehan and Song, 2016, Schrider and Kern, 2018, Flagel et al., 2019, Huang et al., 2024]. Another benefit is that ML models come in many structures that can accept and combine different types of input data. One type of ML model is a dense neural network (DNN). Similar to ABC, DNNs can make predictions about an outcome variable from a vector of summary statistics, but the underlying architecture relies on individual computational units or “neurons” whose outputs are weighted and combined to give the final outcome variable [Sze et al., 2017]. For example, one DNN-based tool called Locator can predict the geographic location of a sample from mostly just vectors of derived allele counts [Battey et al., 2020]. Another type of ML model, convolutional neural networks (CNNs), are specifically designed to process images. Since images are essentially matrices of color values, CNNs are a natural choice for analyzing genotype matrices. CNNs have proven useful for a variety of population genetics tasks, from the prediction of selective sweeps [Kern and Schrider, 2018, Torada et al., 2019, Nguembang Fadja et al., 2021, Whitehouse and Schrider, 2023, van den Belt et al., 2024], to adaptive introgression sites [Gower et al., 2021], dispersal rates [Smith et al., 2023, Smith and Kern, 2023], admixture [Blischak et al., 2021, Rincón Barrado et al., 2024], recombination rates [Flagel et al., 2019], and coalescence times [Nait Saada et al., 2023]. CNNs and DNNs specific to population genetics have even been packaged into standardized tools that can be retrained on broad classes of problems [Sanchez et al., 2022, Zhao et al., 2023]. Altogether, the broad range of inputs that ML models can accept and their lack of reliance on presupposed analytical models creates broad hope that ML can uncover new patterns that summary statistics previously missed (e.g. Mondal et al. [2019], Hahn and Mishra [2025]).

Our core question is whether ML models, with their ability to learn features directly from data, can uncover any new patterns that are useful for predicting *t_f_* that are not already captured by known summary statistics. There are not many examples yet of ML models uncovering brand new patterns in population genetics, but one notable example is that CNNs uncovered a new pattern that recombination rate variation leaves in the site frequency spectrum [Hahn and Mishra, 2025]. Despite this success, CNNs often perform similar to summary statistics for inferring demographic history [Sanchez et al., 2021] or inferring sweeps [Whitehouse and Schrider, 2023]. CNNs will often rely heavily on haplotype structure [Cecil and Sugden, 2023, Tran et al., 2025] which is already quantified by several summary statistics. Thus, whether CNNs can uncover any new signals for differentiating *t_f_*and *t_a_*is an open question. We directly compared the performance of a common CNN architecture [Flagel et al., 2019] and ABC for predicting *t_f_* on simulated data. As an intermediate, we also included DNNs, which are trained on summary statistics (like ABC) but have a neural network architecture (like CNNs) and a CNN model directly fed summary statistics (CNN+DNN). We tested a range of demographic models of panmictic populations and focused on unphased genotype data, which is more readily available in non-model systems and often performs comparably to phased data [Kern and Schrider, 2018, Harris et al., 2018, Mughal and DeGiorgio, 2019, Gower et al., 2021, Arnab et al., 2023]. The hypothesis was that the CNN we tested could more accurately predict *t_f_*than ABC or DNNs, but this turned out to be false.

## Materials and Methods

Our entire analysis is available as a snakemake (v9.33.0) workflow on github (https://github.com/milesroberts-123/slim-sweep-cnn) [Köster and Rahmann, 2012, Mölder et al., 2021]. The workflow is formatted according to standardized snakemake criteria [Mölder et al., 2021] and includes all of the scripts used to simulate data, fit models, and generate figures as well as the configuration files specifying the exact software versions used for each step. The software environments are also packaged into a container on dockerhub so that the analysis can be replicated without installing any software (https://hub.docker.com/r/milesroberts/slim-sweep-cnn). All statistical analysis and plotting are recorded in an R notebook (V4.4.0).

### Simulations

We simulated selective sweeps in SLiM (v4.0.1, Haller and Messer [2019]) with parameters drawn from the distributions listed in Table 1. Similar to previous works, we drew parameters from log-uniform distributions, such as for the selection coefficient (*s*), so that the bulk of simulations did not reflect overly strong selection [Flagel et al., 2019, Torada et al., 2019, Whitehouse and Schrider, 2023]. We performed two separate batches of simulations. **Batch 1** consisted of approximately 200,000 simulations where *t_a_* was allowed to be up to 10,000 generations. Based on these results, we performed an additional 50,000 simulations (**Batch 2**) where we bounded *t_a_* to be 1000 generations or less, as this was the threshold below which we stopped observing non-identifiability (Figure 1). We will first describe the **Batch 1** simulation procedure and then how **Batch 2** built upon **Batch 1**.

**Table 1.** Sampling distributions for key simulation parameters for the five different demographic scenarios included in this study. . U is the uniform distribution and LU is the log uniform distribution, given as (minimum, maximum). Parameters that are identical across all five demographic scenarios are shown once. For *t_a_*, the distributions for **Batch 1** and **Batch 2** simulations are marked with a 1, and 2 respectively.

| Parameter | Constant | Growth | Decay | Cycling | Chaotic |
|---|---|---|---|---|---|
| *N<sub>A</sub>* | U(1000, 10,000) |  |  |  |  |
| *s* | LU(log<sub>10</sub>(1/*N<sub>A</sub>*), 0) |  |  |  |  |
| *h* | U(0, 1) |  |  |  |  |
| *μ* | LU(−8.5, −7.5) |  |  |  |  |
| *R* | LU(−9, −7) |  |  |  |  |
| *t<sub>a</sub>* | **Batch 1**: LU(0, 4); **Batch 2**: LU(0, 0.1*N<sub>A</sub>*) |  |  |  |  |
| *r* | 0 | U(0, 0.5) | U(0, 0.5) | U(2, √6) | U(√6, 3) |
| *K* | *N<sub>A</sub>* | *N<sub>A</sub>* × U(1.01, 2) | *N<sub>A</sub>* × U(0.5, 0.99) | *N<sub>A</sub>* × U(0.8, 1.2) | *N<sub>A</sub>* × U(0.8, 1.2) |

In **Batch 1**, each simulation initialized an ancestral diploid population of size *N_A_*individuals and a chromosome (*L*) of 100 Kb, then burned in for 10*N_A_* generations with mutation rate *µ* and recombination rate *R*. After burn-in, a sweep mutation is introduced to one random chromosome at the middle base of the simulated genome with selection coefficient *s* and dominance coefficient *h*. The population then also has a demography determined by the logistic map:

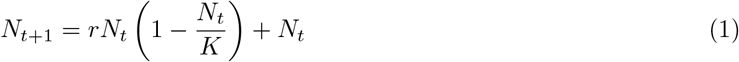

where *N_t_* gives the population size at generation *t* and *N*_0_ = *N_A_*. This equation famously has the following special properties [Phatak and Rao, 1995]:

- *r* = 0, gives a constantly sized population.
- 0 *< r <* 2 and *N*_0_ *< K* gives a population that grows until stabilizing at *K*.
- 0 *< r <* 2 and *N*_0_ *> K* gives a population that decays until stabilizing at *K*.
- 2 *< r <* 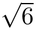 gives a population that cycles between two different sizes.
- 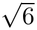 *< r <* 3 gives a population that changes chaotically. In practice, since our population size is a discrete number of individuals, the population can have a long cycle if it happens to return to *N_A_*.

From now on, we will refer to these respective demographies with the terms “constant”, “growth”, “decay”, “cycling”, or “chaotic”, respectively. For growth or decay demographies, we restrict *r* to be 0 *< r <* 0.5 to simulate gradual, rather than instantaneous, population size changes. Because forward simulations are computationally intractable for large populations, we also restrict *K* to be within 1 % - 50 % of *N_A_* (Table 1). For each simulation, we then tracked how long it takes for the sweep mutation to go from frequency 1*/N* to 1 (*t_f_* ). All simulations were conditioned on fixation: if a beneficial mutation drifted out of the population instead of sweeping to high frequency, we restarted the simulation to its post-burn-in state, initialized a new random seed, and re-introduced the beneficial mutation. . After fixation, we allow the simulation to run for an additional *t_a_* generations (sampled from a random distribution Table 1) before taking a random sample of 128 individuals (*n*) from the population, a similar or larger sample size than other sweep-related CNNs [Flagel et al., 2019, Torada et al., 2019, Whitehouse and Schrider, 2023]. Note that *t_a_* is directly sampled from a probability distribution while *t_f_* is not: the exact value of *t_f_* is a random variable whose distribution is a function of other random variables (mostly *N* , *h*, and *s*, Table 1). If a simulation was restarted 1000 times and still did not result in a complete sweep, then we omitted that simulation from downstream analyses.

From the set of all successfully completed simulations with a given demographic pattern, we then randomly downsampled the simulations to create a uniform distribution of log_10_(*t_f_* ). Our downsampling procedure involved taking the initial log_10_(*t_f_* ) distribution, dividing it up into bins of length 0.1 (i.e. 1/10th an order of magnitude), collecting the bin heights, and randomly sampling simulations in each bin (without replacement) until all the bins had the same height as the shortest bin. Our goal with this procedure was to include simulations with fixation times spanning approximately 50 - 20,000 generations while still retaining *>* 11, 000 simulations per demographic scenario. After downsampling, we randomly partitioned the remaining simulations to either the training dataset (80 %), the validation dataset (10 %) or the testing dataset (10 %). Finally, we measured the performance of each model according to the Pearson correlation coefficient between the true and predicted log_10_(*t_f_* ) values and derived confidence intervals for *r* using the cor.test() function in R.

After establishing an initial baseline for the identifiability of *t_f_* and *t_a_* (**Batch 1**), we performed an additional round of simulations (**Batch 2**) with the following adjustments. First, we capped *t_a_* at 0.1*N_A_* (about 1000 generations in the largest simulations), the threshold above which old, fast sweeps left similar signatures as slow, old sweeps (Figure S1C). Second, we redefined *t_f_* as the number of generations to go from a frequency of 0.1 to 0.9, rather than the number of generations to go from frequency of 1*/N* to 1, because the duration of low and high frequency is strongly influenced by drift [Charlesworth, 2020]. Third, instead of a neutral burn-in, we simulated only the sweep mutation’s allele frequency trajectory in SLiM and outputted the result as a tree sequence. We then used pyslim (v1.1.0) to recapitate the tree sequence and then msprime (v1.4.0) to add linked neutral mutations onto the tree. Given that this decreased the runtime of simulations compared to **Batch 1**, we increased the number of sweep attempts from 1000 to 10,000, allowing us to include weaker sweeps in the analysis. We next increased the chromosome size from 100 Kb to 5 Mb, to allow for the exploration of larger context windows around the sweep. With the larger chromosome, we trained models on a window of 128 or 256 SNPs closest to the sweep site. Before train-test partitioning, we did not downsample *t_f_* to a uniform distribution as this seemed to not have any effect on the results and this procedure required massive amounts of oversimulation; the total number of simulations was always capped by the number of simulations in the smallest frequency bin. However, our total dataset size per demographic scenario remained the same between **Batch 1** and **Batch 2** (*∼*11,000). For **Batch 2** simulations, we also checked observations of *t_f_*against a theoretical expectation from [Ewens, 2004]:

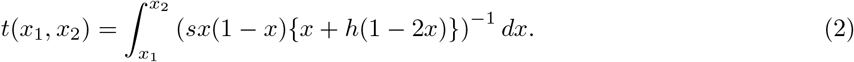

where *t*(*x*_1_*, x*_2_) is the time for a mutation to go from frequency *x*_1_ to *x*_2_ given that it will eventually reach

*x*_2_. We solved this integral with R’s built-in integrate functions and used the simulation parameters as *s* and *h*. Note that this formula does not include *N* , but it is a reasonable approximation given that our simulations had mostly high values of *N* (Figure S7).

#### Selective sweep statistics

For each simulation, we used the VCF file outputted by SLiM and a custom scikit-allel (v 1.3.8) Python script to calculate a suite of selective sweep summary statistics in the window of up to 128 (or 256) variants closest to the site of the sweep, ignoring phase information (similar to previous work [Kern and Schrider, 2018]). We refer to the multilocus diploid genotypes in these unphased data as diplotypes, following Kern and Schrider [2018], and to the unphased analogs of haplotype statistics calculated from them as diplotype statistics. Our statistics included the number of segregating sites (*S*, up to a max of 128 or 256), nucleotide diversity (*π*), Watterson’s theta (*θ_W_* ), Tajima’s D [Tajima, 1989], variance in Tajima’s D, number of diplotypes [Kern and Schrider, 2018], diplotype analogs of Garud’s haplotype statistics (h1, h2, h12, h123, h2h1) [Kern and Schrider, 2018], variance; skew; and kurtosis in the distribution of genotype mismatches (i.e. the *g_kl_* statistic from Kern and Schrider [2018]), average genotypic correlation between pairs of SNPs (Rogers and Huff’s *R*^2^, Rogers and Huff [2009]), Kim’s *ω* [Kim and Stephan, 2002], and unphased Messer’s hscan [Schlamp et al., 2016]. We also span-normalized the unphased Messer’s hscan statistic by dividing the statistic by the length of the 128/256 variant window in bases to get a value between 0 and 1, corresponding to a low and high hscan statistic respectively (Figure S8).

#### Partial *R*^2^ calculations

To investigate the benefit of adding more selective sweep statistics to explain variation in *t_f_* + *t_a_*, we calculated partial *R*^2^ values. These values can be interpreted as the proportion of additional variation *t_f_* + *t_a_* that is explained by one statistic compared to the other 16 statistics.

The procedure to calculate partial *R*^2^ is to first fit an ordinary least squares linear model, which we will call the full model:

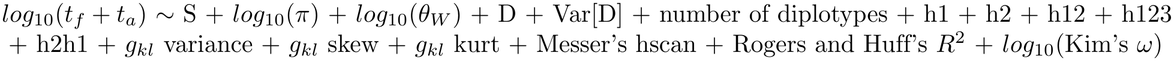

Then, fit a second model with one term removed. If we call the former model the full model (*M* ) and the latter model the reduced model (*m*), then the partial *R*^2^ of the dropped term is:

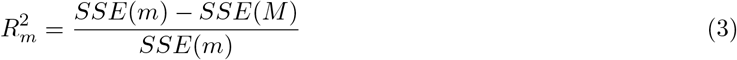

where *SSE* is the sum of squared errors.

#### Image-based sweep representation

In addition to calculating sweep statistics, we also converted the VCF file output from each simulation into an image representation using a custom R script with ggplot [Wickham, 2016], ggnewscale [Campitelli, 2022], and cowplot [Wilke, 2020]. This image representation is similar to [Flagel et al., 2019] but with some modifications for unphased genotype data. Each image was a gray-scale matrix with *n* rows corresponding to the sampled individuals and *l* columns corresponding to the *l* SNPs closest to the selective sweep site. To emphasize haplotype structure [Flagel et al., 2019], we clustered the rows according to their manhattan distance using the complete clustering algorithm in stats R package’s hclust function. The color of each element in the matrix could be either black (homozygous genotype 0/0), grey (heterozygous genotype 0/1), or white (homozygous genotype 1/1). We allowed *l* to reach a maximum of 128 or 256 SNPs and padded each image with black columns if there were fewer than 128 or 256 SNPs in the sampled simulated population. Because each column of the image represents a SNP we also created a vector of SNP positions to go along with each image. These SNP positions were min-max normalized such that the SNP in the left-most column of an image was position 0 and the SNP at the right-most column of the image was position 1.

#### Convolutional Neural Networks

We used the image-based representations of genotypes in the 128/256 variant window near the sweep site to train a CNN using keras (v2.11.0, Chollet and others [2015]) and tensorflow (v2.14.0 Martín Abadi et al. [2015]). All activation functions used rectified linear units [Agarap, 2018]. Our CNN architecture was similar to Flagel et al. [2019], with two main branches: one to process a sweep image and another to process the vector of SNP positions associated with the columns of the sweep image. The image processing branch had three convolutional layers, each followed by a pooling and dropout layer. The first convolutional layer had a kernel size of 7 and a stride of 2. The second and third convolutional layers had a kernel size of 3 and a stride of 1. The final dropout layer was flattened and fed into a dense layer. The branch for processing the position information began with an input layer with 128 or 256 neurons (one for each column in the image), followed by a dense layer and a dropout layer. The final dense layer for both the image-processing and position-processing branch were concatenated and fed into a final dense layer with dropout before being fed into a single output neuron.

To determine the proper dropout rates and sizes of the convolutional and dense layers, we performed 60 iterations of Bayesian hyperparameter tuning implemented in keras on each CNN. We chose 60 iterations because a random parameter search with 60 iterations will sample a model in the top 5 % of the performance range with probability 95 % [Bergstra and Bengio, 2012]. Thus, we would expect 60 iterations of a Bayesian hyperparameter approach to perform at least as well. Each iteration involved 2 epochs (for **Batch 1** simulations) or 8 epochs (for **Batch 2** simulations, to allow more time for CNNs to learn) of model fitting to predict *log*_10_(*t_f_* ) in the training simulations and the final performance of the iteration was measured as mean squared error between the predicted and actual *log*_10_(*t_f_* ) values on the validation simulations. During tuning, each convolutional layer could have any multiple of 16 filters between 16 and 128 and any layer could have a dropout rate between 0 and 0.99. All dense layers could have any multiple of 32 neurons between 32 and 512. The first convolutional layer was fixed to have a kernel size of 7 and a stride of 2, while the second and third convolutional layers had a kernel size of 3 and stride of 1. All activation functions were rectified linear unit (ReLU) functions [Banerjee et al., 2019].

After hyperparameter tuning, we then trained a final CNN with the best performing hyperparameters. Each training epoch fit the CNN in batches of 32 images and we randomly flipped the training images either horizontally or vertically to prevent the model from fixating on the exact arrangement of rows and columns. We trained the CNN until validation performance did not improve for 20 epochs. After 20 epochs of no improvements, we reverted the CNN to the configuration with the best performance on the validation data. This early stopping criterion is meant to avoid overfitting the model on the training data [Prechelt, 1998]. Final model performance was assessed on the testing data and we performed 100 iterations of Monte-Carlo sampling [Gal and Ghahramani, 2015] on each testing image and used the mean of the predictions across the Monte-Carlo samples as the point estimate for *t_f_* from the CNN. For **Batch 2** simulations, we also measured the best performing model for the given demographic scenario as the model with lowest root-mean square error (RMSE) on the validation set, calculated as:

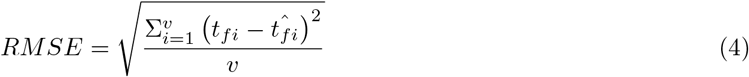

where *v* is the number of simulations in the validation set and *t_f_* and *t*^^^*f* are the true and predicted values of *t_f_* , respectively.

#### Dense Neural Networks

We also constructed dense neural networks (DNNs) that were trained on only summary statistics. Our DNNs had an input layer of 17 neurons (one for each summary statistic) followed by three dense layers with dropout. Similar to the training of CNNs, we did 60 iterations of hyperparameter tuning using Bayesian hyperparameter optimization, with 2 epochs per iteration for **Batch 1** or 8 epochs per iteration for **Batch 2**. During tuning, each of the dense layers could have a multiple of 8 neurons anywhere from 16 to 512 and a dropout rate anywhere from 0 to 0.99. Final model performance was assessed on the testing data and we performed 100 iterations of Monte-Carlo sampling on each testing example. The mean of the predictions across the Monte-Carlo samples was used as our point estimate for *t_f_* . Finally, we implemented a combined DNN+CNN approach where we combined this DNN architecture with the CNN architecture described above. This approach gave the CNN more pre-computed information on sweep features, potentially helping it focus on discovering small residual sweep features not represented in the summary statistics rather than discovering already known sweep features.

#### Approximate Bayesian Computation

We also performed ABC to estimate *t_f_* using the abc package (v2.2.1, Csilĺery et al. [2012]) in R [R Core Team, 2022]. Just like tuning hyperparameters of CNNs, performing ABC requires making several choices, including the choice of regression method, the tolerance level, and the type of point estimate used (i.e., mean, median, or mode of the posterior distribution). Thus, similar to ML, it is typically desirable to try many hyperparameter choices and identify a combination of hyperparameters with high performance to use for final predictions. For each demographic scenario, we tried 63 different configurations of ABC (similar to doing 60 iterations of hyperparameter tuning for CNNs and DNNs), including 3 different regression methods (rejection, ridge regression, local linear regression), 7 different tolerance values (0.025, 0.05, 0.1, 0.15, 0.2, 0.25, 0.3), and 3 different posterior distribution point estimates (mean, median, or mode). For each model configuration, we predicted *t_f_* for each simulation in the validation set using the training set as the baseline. We quantified the performance of the model as the Pearson correlation between the model’s predictions and the true value, and we also calculated the RMSE of the predictions for **Batch 2** simulations. For the best performing ABC model, we then measured final performance by making predictions for the testing dataset, using the training dataset as the background.

#### Simple Linear Regression

As a baseline for comparison to the more complex CNN, DNN, and ABC models, we also included some ordinary linear regressions of individual summary statistics vs *log*10(*t_f_* ). For each demographic scenario, we used the same training vs testing partitions of simulations sets as we did for the other models above. We fit simulations in each training partition to a linear model of either *log*10(*t_f_* ) *∼ log*10(*π*) or *log*10(*t_f_* ) *∼* hscan using the lm() function in base R. We then predicted *log*10(*t_f_* ) for held out simulations in the testing sets of simulations for each demographic scenario using the predict() function in base R. We measured model performance as the RMSE of *log*10(*t_f_* ) predictions in the testing set (the same as for all the other models above).

## Results

### Simulations and statistics reflect known challenges disentangling *t_f_* and *t_a_*

We performed a total of 250,000 SLiM simulations across 5 different demographic scenarios (50,000 simulations per scenario). In **Batch 1** simulations, 79% of simulations successfully produced hard sweeps with valid values of all 17 selective sweep statistics. In **Batch 2** simulations, *>* 97% of simulations produced complete hard sweeps with non-NA values for all sweep statistics in all demographic scenarios. For each of these simulations, we recorded *t_f_*and *t_a_*for the sweep (Figure 1A). In the constant demography scenario, *π* monotonically increased with *t_f_* + *t_a_* (Figure 1B). If *t_a_* was unbounded, sweeps that were old (*t_a_ >* 1000) and fast (*t_f_ <* 100) generally had similar values of sweep statistics compared to sweeps that were younger (*t_a_ <* 1000) and slower (*t_f_ >* 100, Figures S1C, S2-S5, S4, S3). But when we capped *t_a_*, summary statistics better distinguished the *t_f_*continuum (Figure 1C). Overall, the selective sweep statistics explained at least 78.7 % of variation in *t_f_* based on a simple linear model, and the amount of explained variation increased to *>* 80% if the length of SNP context surrounding the sweep site was increased from 128 to 256 (Table S1).

We used partial *R*^2^ values to better understand the importance of each statistic for explaining variation in *t_f_* + *t_a_*. First, we investigated correlations among statistics. The statistics generally fell into three classes: (1) site frequency spectra statistics, (2) haplotype frequency statistics, and (3) linkage disequilibrium–based statistics. Statistics within the same class were generally positively correlated with each other since they measure fundamentally similar features (Figures 2A, 2B, S6). However, SFS statistics were generally negatively correlated with the haplotype frequency statistics since more mutations in a region generally correspond to shorter haplotypes (or, in our case, diplotypes), all else being equal. Interestingly, the three variants of the *g_kl_* statistic, which are simply three different moments of the *g_kl_* distribution, fell into separate correlation modules: *g_kl_* variance correlated with site-frequency spectra statistics, *g_kl_* kurtosis correlated very little with other statistics, and *g_kl_* skew correlated with haplotype statistics. Similarly, Tajima’s *D* and Var[*D*] were only slightly correlated with each other. Rogers and Huff’s *R*^2^ and Kim’s *ω* were generally not correlated with any of the other statistics, except for being slightly positively correlated to each other (Figures 2A, 2B), potentially reflecting that these were the only statistics quantifying linkage disequilibrium. Across the 5 different demographic models, Messer’s Hscan generally had the highest partial *R*^2^ with the exception of the chaos demographic scenario (Figure S6C, S6F). Tajima’s D, *π*, Kim’s *ω*, and Rogers and Huff’s *R*^2^ generally had the highest partial *R*^2^, though most partial *R*^2^ values were *<* 0.07, suggesting that there is overlap in the information provided by individual summary statistics (Figures 2C, 2D, S6D-S6F).

**Figure 2.**
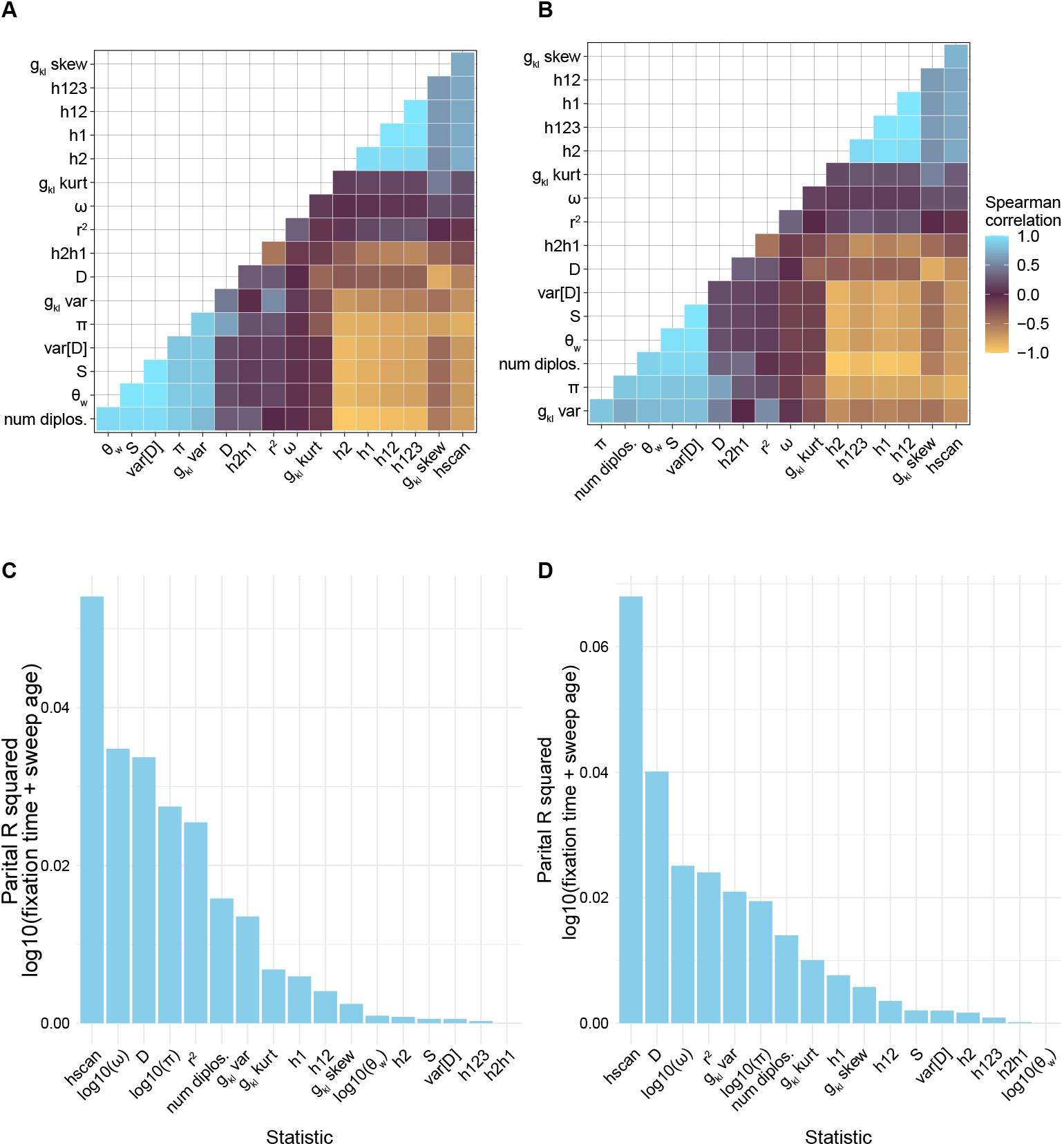
For each simulation we calculated 17 summary statistics. (A) and (B) show the pairwise Spearman correlations between statistics in the final stratified dataset used for training in the constant and growth scenarios, respectively, for **Batch 1** simulations. (C) and (D) show the partial *R*^2^ of each statistic for explaining variation in *log*_10_(*t_f_* + *t_a_*) calculated with Equation 3 in the constant and growth scenarios, respectively, for **Batch 1** simulations. These statistics were nucleotide diversity (*π*), the number of segregating sites (S) up to a max of 128, Watterson’s theta (*θ_W_* ), Tajima’s D, variance in Tajima’s D (*V ar*[*D*]), mean LD (*r*^2^), number of diplotypes (num diplos.), distribution of genotype mismatches (*g_kl_* var., *g_kl_* skew, *g_kl_* kurt), unphased equivalents of Garud’s haplotype frequency statistics (h1, h2, h12, h123, h2h1), Kim’s *ω*, and Messer’s Hscan.

### Convolutional neural networks vs summary statistics for disentangling *t_f_* and *t_a_*

For each of the 5 demographic scenarios, we built ABC, DNN, and CNN models to predict *t_f_* . The Pearson correlation between the true value of *t_f_* and the predicted value of *t_f_* for the best performing version of each model was generally *>* 0.7 (Figure S9, S10, S11-S13). By this measure of performance, the three types of models were not significantly different for most of the demographic scenarios tested (e.g. 95 % confidence intervals for correlation between true and predicted *t_f_* values in constant demography: CNN = [0.705, 0.750], DNN = [0.719, 0.762], ABC = [0.731, 0.773]; Figure S9) with the exception of the cycling demography where the CNN performed worst (r = 0.656 (CNN) vs 0.728 (DNN) vs 0.691 (ABC), Figure S12). These **Batch 1** simulations revealed an obvious identifiability threshold for *t_a_*: sweep ages *>* 1000 generations (or 0.1*N_A_* for our larger simulations) caused difficulty in estimating *t_f_*across all models tested. Models were incorrectly predicting that sweeps with high *t_a_* had long *t_f_* (Figure S9). The one exception was the growth demography scenario where the CNN and DNN appeared to confuse old, fast and slow, young sweeps less often (Figure S10), but failed to achieve statistically better performance than ABC (*r* =CNN: 0.75, DNN: 0.781, ABC: 0.77).

We further investigated the identifiability of *t_f_* vs *t_a_* with another round of simulation (**Batch 2**, see the Simulations subsection for a full list of adjustments). The main change in **Batch 2** was capping *t_a_* at 0.1*N_A_* to limit confounding between *t_f_* and *t_a_*. We also built simple linear models to predict *t_f_* from the *π* in the sweep region, achieving an RMSE of 0.582 and *r* = 0.799 in the constant demographic scenario (Figures 3 and S14). All other models outperformed these simple linear models (Figures S15-S19). All models also had greater prediction error as *s* got weaker, but prediction error did not correlate with *t_a_* (Figures S24-S33). The CNN often underperformed slightly compared to ABC and DNN in terms of RMSE (Figure S15A-S19A; Figure 4A; Table S2) except for the cycling demography scenario where the CNN improved over DNN (Figure S22B,C). We next created a combined CNN+DNN model, where the summary statistic information is incorporated into the CNN output via three layers of neurons. This improved *t_f_* predictions, in terms of RMSE, in several cases (Figure S16, S17, S18). In two cases, the CNN+DNN model outperformed the ABC (Figure S16) or DNN model (Figure S17). Finally, we increased the window to 256 SNPs and re-fit all of these models (simple linear regression, ABC, DNN, CNN, CNN+DNN). The CNN+DNN combination only performed better than CNN in the decay scenario (Figure S21) but otherwise never outperformed DNN or ABC (Figure 4, S20-S23).

**Figure 3.**
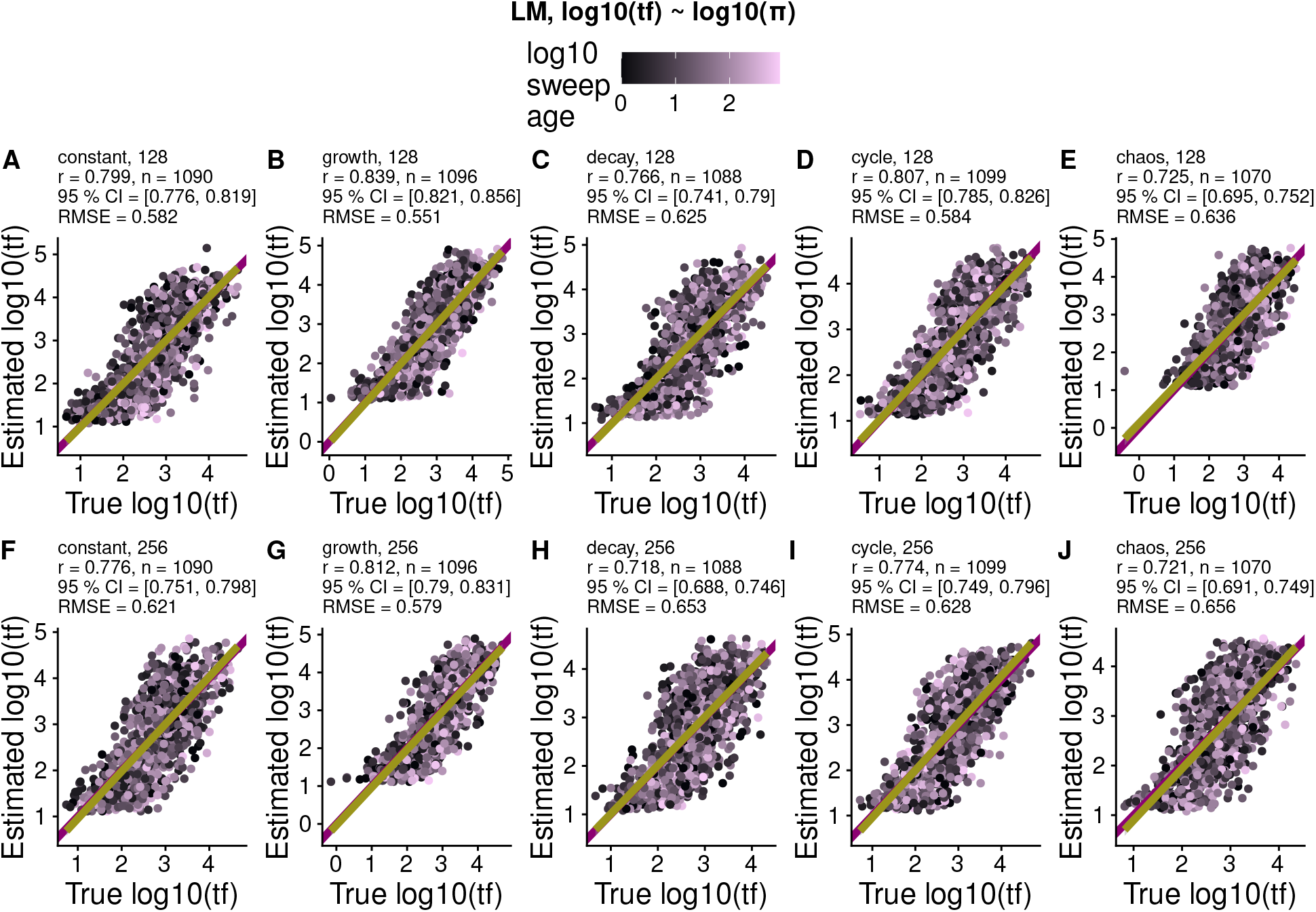
A simple linear model predicting. *t_f_* **from** *π* **performs poorly relative to the other models.** Predicted vs. true log_10_(*t_f_* ) for a linear model fit to log_10_(*π*) within (A-E) 128-SNP and (F-J) 256-SNP windows, for the constant (A, F), growth (B, G), decay (C, H), cycling (D, I), and chaotic (E, J) demographic scenarios. For each panel we list the Pearson correlation coefficient between predicted and true values (*r*), the number of test set examples (*n*), the 95 % confidence interval for *r*, and the root mean square error of the predictions from ground truth (RMSE). All points are colored according to the *t_a_* of the sweep in generations (log10 transformed).

**Figure 4.**
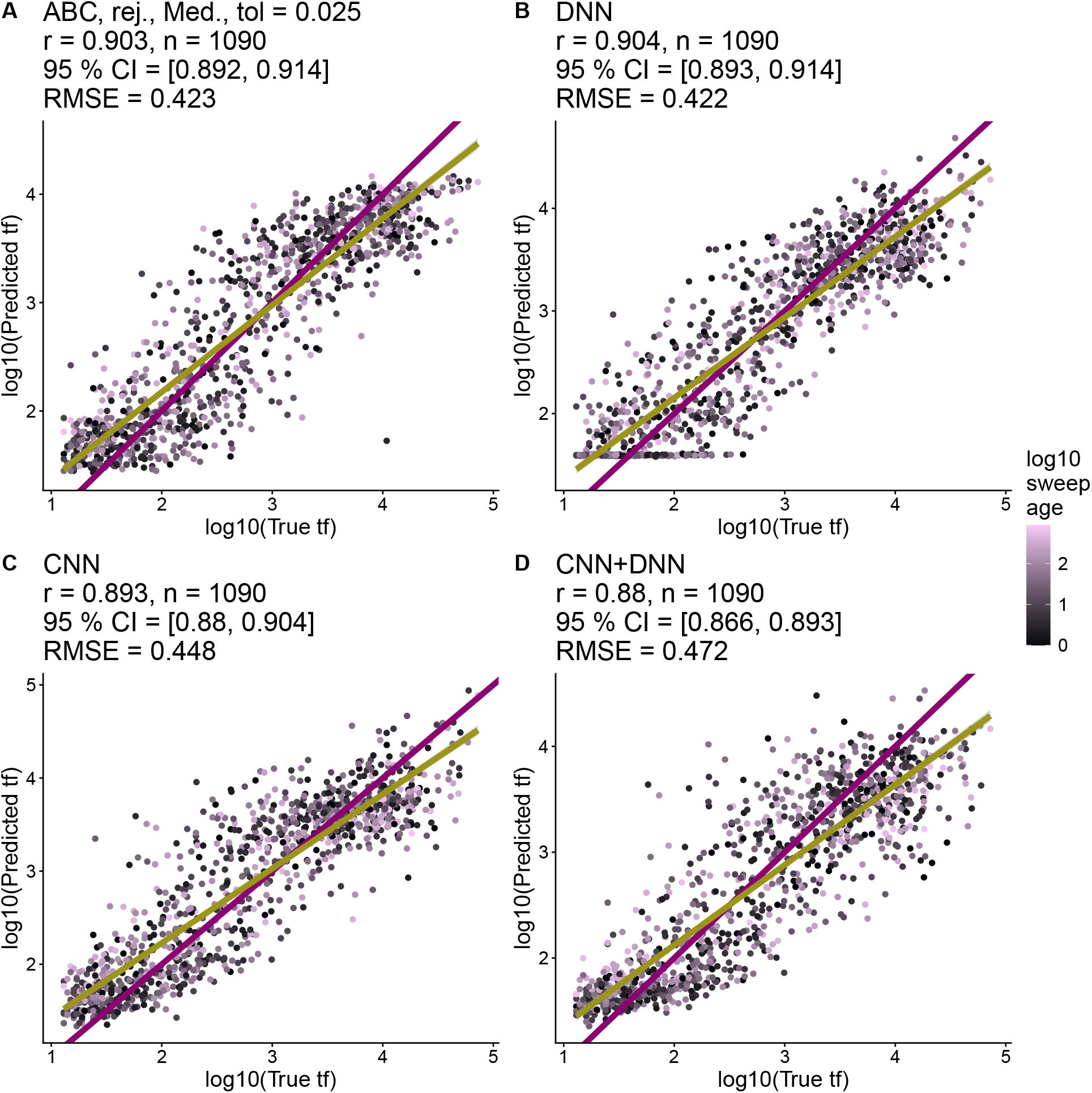
Performance of CNN, DNN, and ABC models in Batch 2 simulations, 256 SNP window, constant demographic scenario. Performance of best (A) ABC, (B) DNN, (C) CNN, and (D) CNN+DNN models at predicting *t_f_* under a constant demographic scenario. The best performing ABC model used the rejection method (rej.), a median posterior estimate (Med.) and a tolerance value of 0.025 (tol). For each model, we list Pearson correlation coefficients between predicted and true values (*r*), the number of test set examples (*n*), the 95 % confidence intervals for *r*, the p-value for testing if *r* is different from 0 (*p*), and the root mean square error of the predictions from ground truth (RMSE). The purple line is the 1-1 reference line where predictions and truth are equal. The yellow line is a least squares regression line. All points are colored according to the *t_a_*of the sweep in generations (log10 transformed).

## Discussion

We simulated approximately 250,000 selective sweeps across a range of parameters and demographic scenarios, then used those simulations to train a simple linear model, a CNN, a DNN, a combined CNN+DNN model, and an ABC model to predict *t_f_* . Our hypothesis was that the CNN could potentially identify new signatures and better predict *t_f_* compared to methods based purely on summary statistics. Our simulations reflected a known challenge in distinguishing *t_f_* from *t_a_*: sweeps with high *t_f_* and low *t_a_* (i.e. slow, young sweeps) leave similar signatures as sweeps with low *t_f_* and high *t_a_* (i.e. fast, old sweeps, Figure 1C). Similar to previous studies on estimating coalescent times [Brandt et al., 2022], there is a general bias where models overestimate low values of *t_f_* and underestimate high values of *t_f_* (Figure S9, S10, S11-S13). Each of the 17 selective sweep statistics we calculated explained additional variation in *t_f_* + *t_a_* (Figure S6). In the end, our models trained purely on summary statistics (DNN and ABC) performed very similarly to our specific CNN architecture trained on raw genotype matrices (Figure S9, S10, S11, S13).

Surprisingly, our CNN performed significantly worse than the DNNs trained on summary statistics in the cyclic demography scenario (*r* = 0.656 vs 0.728, Figure S12), suggesting that summary statistics provided useful information that the tested CNN was unable to learn. We then attempted to feed summary statistics directly into the CNN [Lauterbur et al., 2023], allowing the CNN to skip needing to learn known selection signals and focus on discovering any remaining residual signal. While this often improved performance compared to CNN, this combined CNN+DNN model never outperformed both ABC and DNN alone in the same scenario (Figure S15D, S16D-S23D). This suggests that the tested CNN can struggle to distinguish spurious from true signals under certain demographic scenarios. Increasing the number of training simulations could improve performance on these demographies. However, even after downsampling our simulations to have a uniform distribution of *t_f_* , we had *>*11,000 simulations for each demographic scenario, which is on par with many previous CNN based models in population genetics [Kern and Schrider, 2018, Flagel et al., 2019]. This suggests that any undiscovered signatures to discriminate *t_f_* and *t_a_* in hard sweeps (if they exist) could be very subtle and require many more simulations and training effort to detect.

There is promise that applications of ML to population genetics will uncover novel signatures of evolutionary processes [Hahn and Mishra, 2025]. However, systematic searches for new signals with ML are rare in the literature. At least one previous study indicated that CNNs trained on genotype matrices to identify sweeps largely rely on known signals of haplotype structure and will recapitulate known haplotype statistics [Cecil and Sugden, 2023, Tran et al., 2025]. This makes sense because measures of haplotype length seem particularly powerful for explaining variation in *t_f_*and *t_a_*(Figure S6). However, if the goal is discovery of new signals, one could maybe penalize models that begin to reproduce known summary statistics, forcing them to look for new patterns. This approach is somewhat similar to new methods that improve generalization by penalizing models for overfitting on simulated data [Mo and Siepel, 2023], but has not yet been attempted in the literature. Another option is to give CNNs access to more types of data, potentially allowing them to combine information in genotype matrices. For instance, it was recently discovered that sweeps leave signatures in the spatial distribution of genotypes [Rehmann et al., 2025] and such data could also be encoded into images for a CNN. Many CNNs are also trained on phased or polarized genotype matrices [Torada et al., 2019] or time series data [Whitehouse and Schrider, 2023], but such data are often not available in non-model species and phasing does not always appreciably improve the detection of positive selection in ML-based methods [Kern and Schrider, 2018, Harris et al., 2018, Mughal and DeGiorgio, 2019, Gower et al., 2021, Arnab et al., 2023]. We observed an inability to distinguish between strongly selected ancient sweeps and weakly selected young sweeps nearly universally across our parameter space, reflecting similar benchmarks for coalescent time estimation [Brandt et al., 2022]. However, in our **Batch 2** simulations, where *t_a_* was capped at a max of 1000, *t_f_*prediction error was related to *s* rather than *t_a_*(Figure S24 - S33). This result suggests that *t_f_*inferences are robust to a certain threshold of *t_a_*: approximately 1000 generations in our simulations. However, beyond this threshold, any method will struggle to recover *t_f_* as it becomes fundamentally confounded with *t_a_*. If this pattern reflects a fundamental limitation of single-time point population genetic observations, then we might expect other types of timing estimates to be similarly affected, such as timing of deleterious sweeps [Johri et al., 2021] and coalescent events . However, we cannot eliminate the possibility that CNNs could discover new signatures to discriminate *t_f_*and *t_a_*if given additional information or trained on new demographic scenarios not explored here . We attempted to explore a wide range of demographic space, and our generalizable workflow can easily be extended to new demographies, classes of sweeps, or additional configurations of genotype matrices. Perhaps greater focus on time series population genetic analyses will help push the bounds of *t_a_* further.

## Data availability

Our entire analysis is available as a snakemake workflow, packaged here: https://github.com/milesroberts-123/slim-sweep-cnn. All of the software for the workflow is packaged as a container on dockerhub: https://hub.docker.com/r/milesroberts/slim-sweep-cnn. Upon acceptance of publication, we are also prepared to publish our trained models and the image datasets we used for training in the accepting journal’s preferred repository.

## Acknowledgments

We would like to thank Yaniv Brandvain, Jeff Conner, Bob VanBuren, Shinhan Shiu, Addie Thompson, Husain Agha, Sophie Buysse, Nathan Catlin, Maya Wilson Brown, Asia Hightower, and Derek Denney for feedback on early drafts of the manuscript.

## AI Use Disclosure

No generative AI tools were used for the conception, data analysis, or writing for this project. Overleaf’s built-in AI helper was occasionally used to fix L^A^T_E_Xcompilation warnings and errors. We also used GLM5.3-Flash agents through Ollama Cloud to resolve cross-document line number and figure referencing errors in L^A^T_E_Xand minor wording corrections for typos and grammar checking, all of which were human reviewed. The agents did not alter the content of the writing. The full transcript of the agents’ activities is available as a compressed .json on github (https://github.com/milesroberts-123/cnns-time-to-fixation-manuscript). Gemini and Claude Sonnet were occasionally used to search for boilerplate R code to load files and make plots in ggplot, but every line of code was also human-reviewed.

