## Supplemental tables and figures for "Summary Statistics and Approximate Bayesian Computation Are Comparable to a Common Convolutional Neural Network Design for Inferring Fixation Times"

November 16, 2026

### 1 Initial analyses

#### 1.1 Batch 1: Checking for unidentifiability in summary statistics for predicting $t_f$

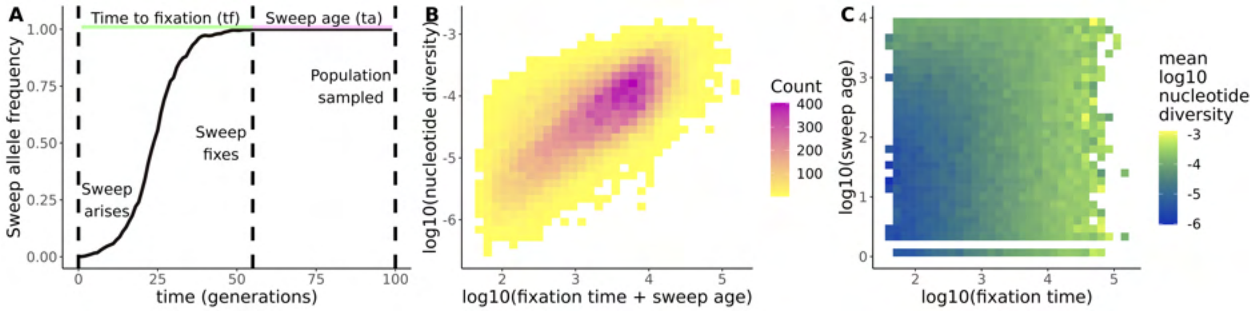

Figure S1: **Separately estimating $t_f$ and $t_a$ from summary statistics leads to unidentifiability.** (A) Example allele frequency trajectory comes from simulating a beneficial allele with selection coefficient 0.5, dominance 0.5, and initial frequency of 1/1000 in a Wright-Fisher population of size 1000. (B) Nucleotide diversity in the sweep region linearly scales with the total lifetime of the sweep: $t_f + t_a$. (C) Young, slow sweeps (high $t_f$ and low $t_a$) have similar nucleotide diversity as old, fast sweeps (low $t_f$ and high $t_a$).

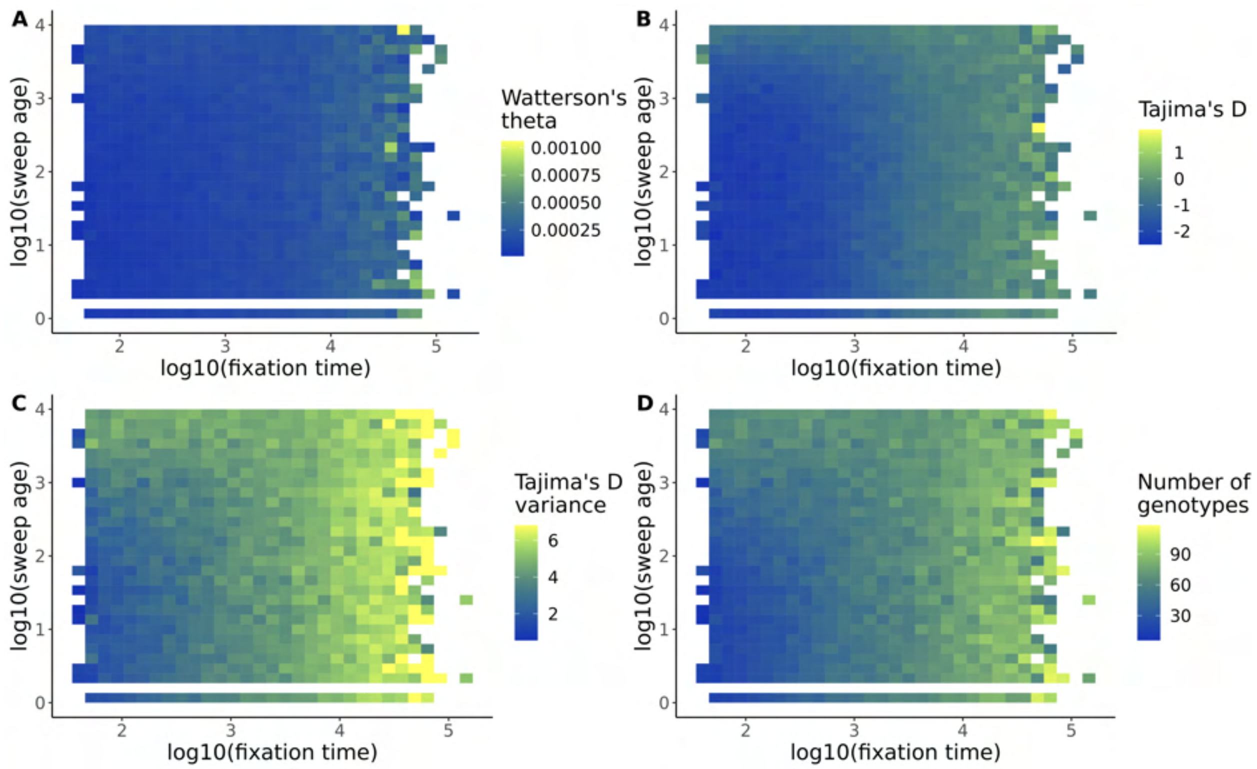

Figure S2: $t_f$ **vs** $t_a$ **for site frequency spectrum summary statistics.** Every data point is a simulated sweep and plots show how (A) Watterson's theta ($\theta_W$), (B) Tajima's D, (C) Tajima's D variance (i.e. the denominator of Tajima's D), and the number of diplotypes in the 128 SNP regions centered around the sweep site vary based on the fixation time and age of the sweep. All simulations were done in constant-sized populations.

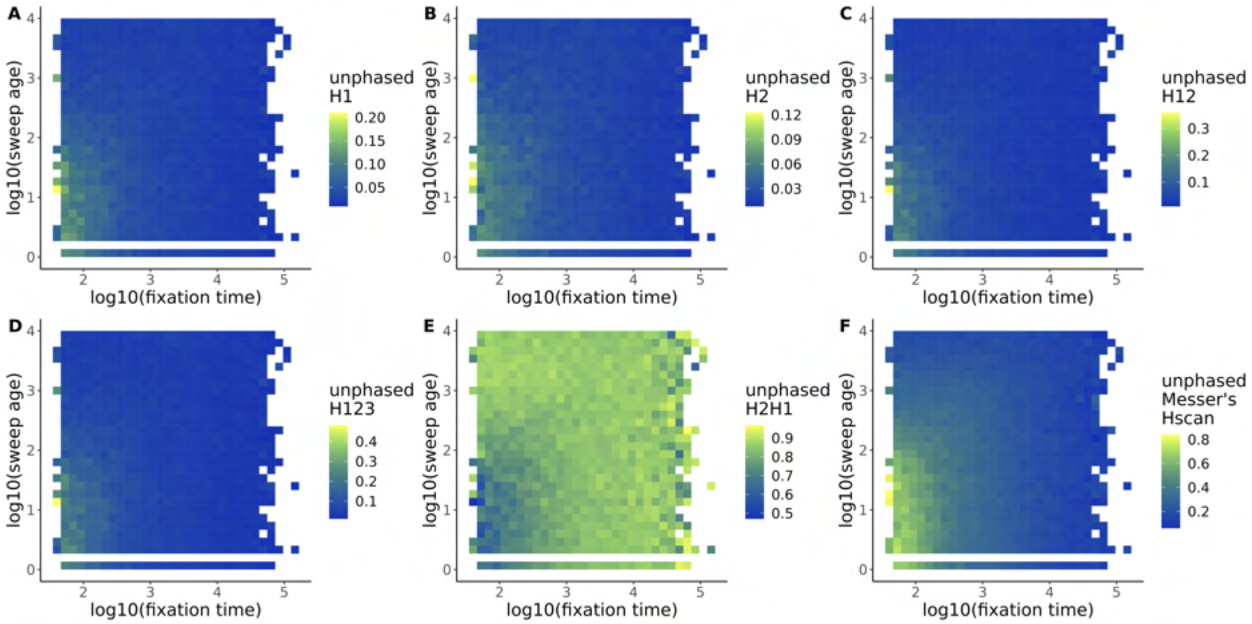

Figure S3: $t_f$ **vs** $t_a$ **for unphased haplotype/diplotype-based summary statistics.** Every data point is a simulated sweep and plots show how (A)-(E) Garud Haplotype statistics (unphased equivalents), and (F) unphased Messer's Hscan vary with the fixation time and age of the sweep. All simulations were done in constant-sized populations.

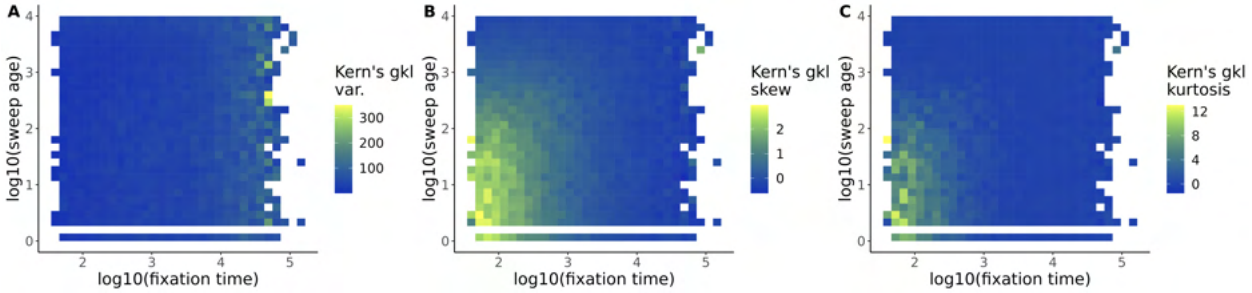

Figure S4: $t_f$ **vs** $t_a$ **for Kern's** $g_{kl}$ **statistics in constant-sized populations.** All simulations were done in constant-sized populations.

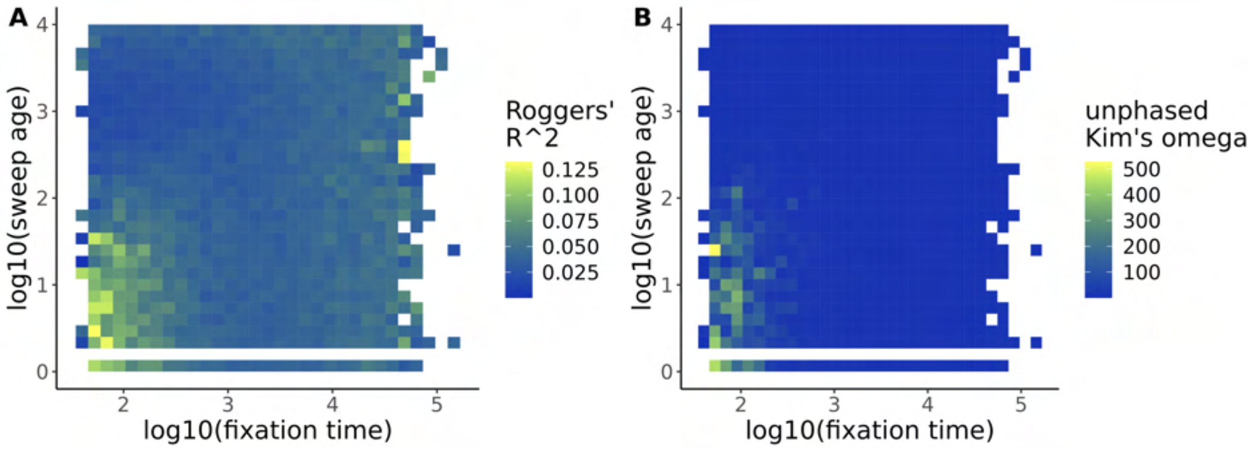

Figure S5: $t_f$ **vs** $t_a$ **linkage disequilibrium-based summary statistics.** Every data point is a simulated sweep and plots show how (A) Rogers' $R^2$ and (B) Kim's $\omega$ vary with the fixation time and age of the sweep. All simulations were done in constant-sized populations.

#### 1.2 Batch 1: Checking that summary statistics add additional information to predicting $t_f$

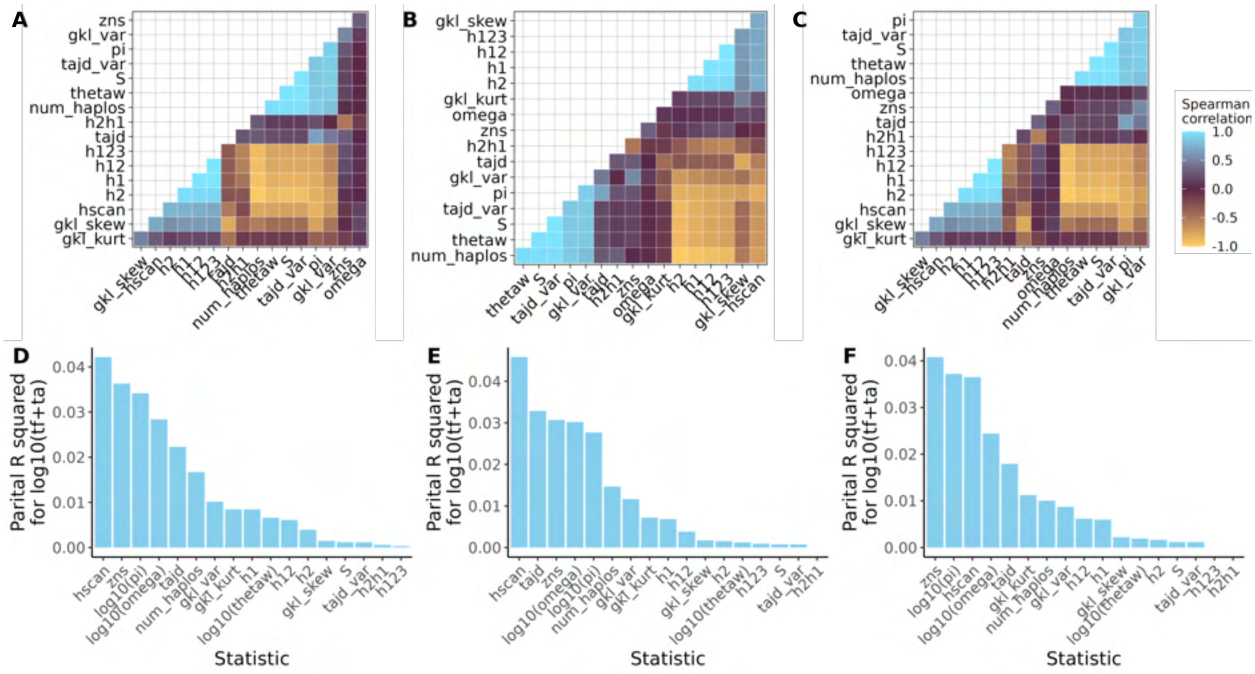

Figure S6: **Relationships between summary statistics and $t_f + t_a$.** (A), (B), and (C) show pairwise Spearman correlations between selective sweep statistics in the decay, cycle, and chaos demography scenario respectively. (D), (E), and (F) show partial-R-squared values for each summary statistic describing amount of additional variation in $t_f + t_a$ explained by each statistic compared to all other statistics. Statistics with lots of extreme values were log-transformed as indicated in the plot before calculating the partial $R^2$ values to improve the fits of the linear models. The statistics include Rogers and Huff's $R^2$ (zns), $\pi$ (pi), Tajima's D (tajd) and its variance (tajd_var), Watterson's theta (thetaw), the number of diplotypes (num_haplos), number of segregating sites (S), and Kim's $\omega$ (omega).

#### 1.3 Batch 2: Checking that theoretical $t_f$ matches expected $t_f$

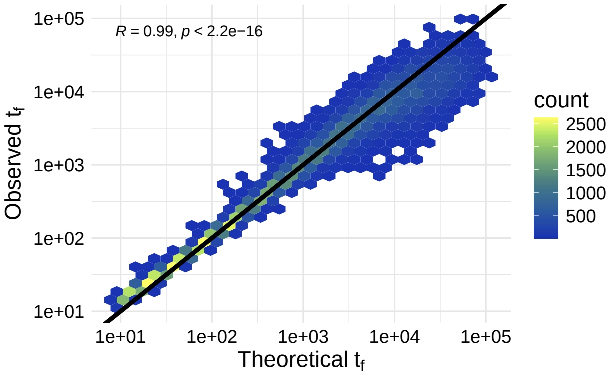

Figure S7: Comparison between theoretical $t_f$ (Equation 3 of the main text) to observed $t_f$ reported from SLiM simulations in the constant demography scenario. The line is a 1-to-1 line where theoretical and observed values would match perfectly. R is the Pearson correlation coefficient and p-value is to test if R is significantly different from zero. To avoid overplotting, points are collapsed into hexbins with counts given by color.

#### 1.4 Batch 2: Checking extent of sweep within analysis window

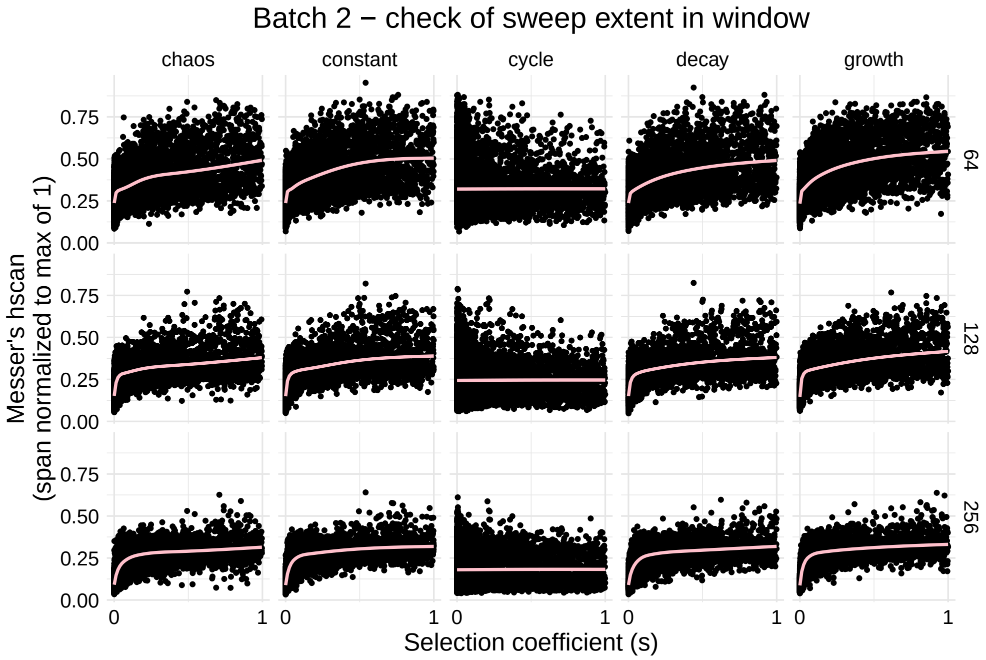

Figure S8: Comparison of Messer's hscan within an analysis window for different window lengths (64, 128, and 256 SNPs for the different rows). A window length of 64 SNPs is plotted for comparison, but these windows were not used to train models. Line is a loess smoothing line.

### 2 Confirmatory analyses

#### 2.1 Batch 1: Comparison of best CNN, DNN, and ABC models in extra demographic scenarios

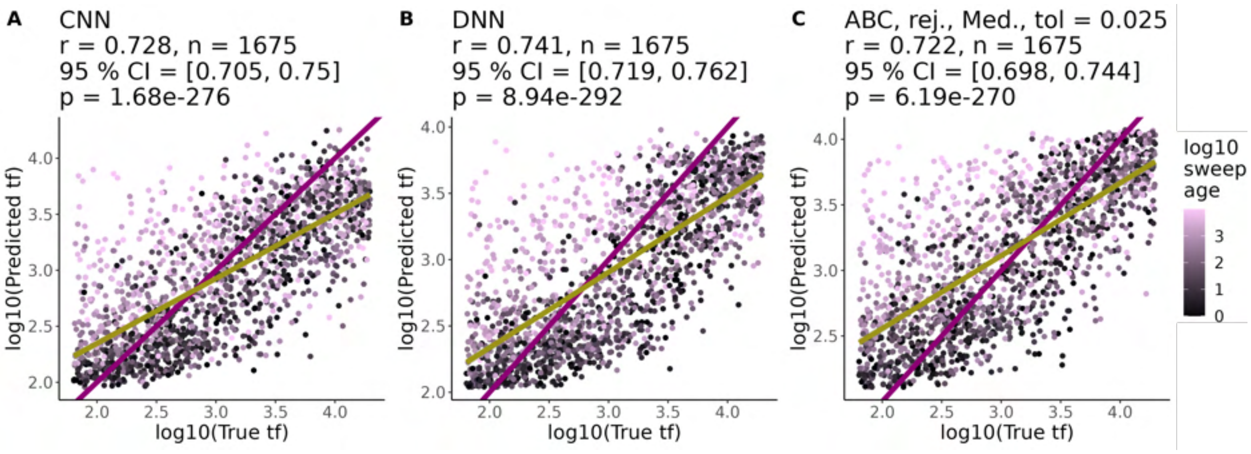

Figure S9: **CNNs perform similarly to DNNs and ABC in a constant-sized population.** Performance of best (A) CNN, (B) DNN, and (C) ABC models at predicting $t_f$ under a constant demographic scenario. The best performing ABC model used the rejection method (rej.), a median posterior estimate (Med.) and a tolerance value of 0.025 (tol). For each model, we list Pearson correlation coefficients between predicted and true values ($r$), the number of test set examples ($n$), the 95 % confidence intervals for $r$, and the p-value for testing if $r$ is different from 0 ($p$). The purple line is the 1-1 reference line where predictions and truth are equal. The yellow line is a least squares regression line. All points are colored according to the $t_a$ of the sweep in generations (log10 transformed).

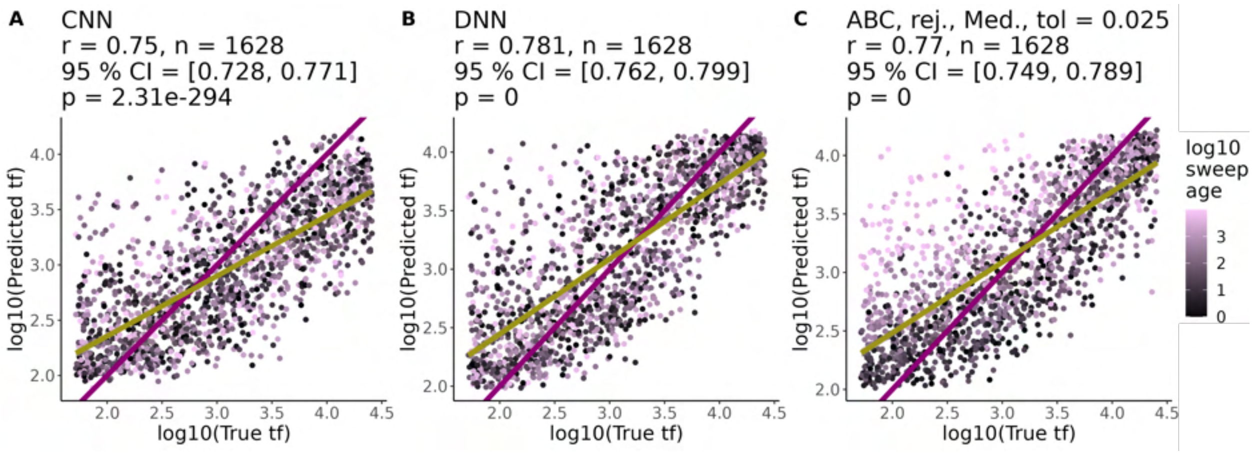

Figure S10: **CNNs, DNNs, and ABC in a growing population.** Performance of best (A) CNN, (B) DNN, and (C) ABC models at predicting $t_f$ under the growth demographic scenario. The best performing ABC model used the rejection method (rej.), a median posterior estimate (Med.) and a tolerance value of 0.025 (tol). For each model, we list Pearson correlation coefficients between predicted and true values ($r$), the number of test set examples ($n$), the 95 % confidence intervals for $r$, and the p-value for testing if $r$ is different from 0 ($p$). The purple line is the 1-1 reference line where predictions and truth are equal. The yellow line is a least squares regression line. All points are colored according to the $t_a$ of the sweep in generations (log10 transformed).

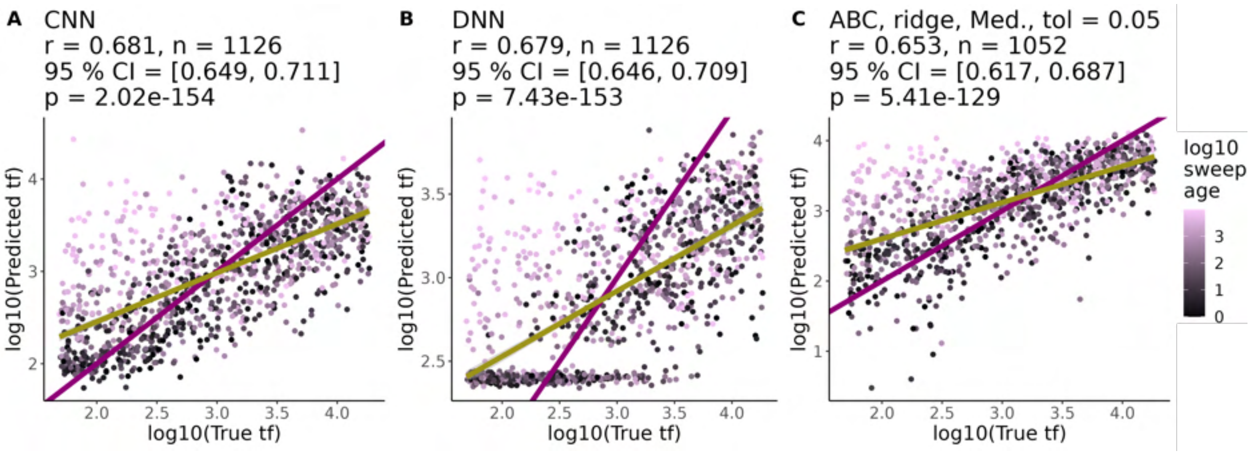

Figure S11: **Comparison of best CNN, DNN, and ABC models at predicting $t_f$ for the decay demography scenario.** Performance of best (A) CNN, (B) DNN, and (C) ABC models at predicting $t_f$ under the decay demographic scenario. For each model, we list Pearson correlation coefficients between predicted and true values ($r$), the number of test set examples ($n$), the 95 % confidence intervals for $r$, and the p-value for testing if $r$ is different from 0 ($p$). The purple line is the 1-1 reference line where predictions and truth are equal. The yellow line is a least squares regression line. For the ABC model, the best performing model was a ridge regression, using the median of the posterior distribution as the point estimate, and a tolerance value of 0.05. All points are colored according to the age of the selective sweep in generations (log10 transformed).

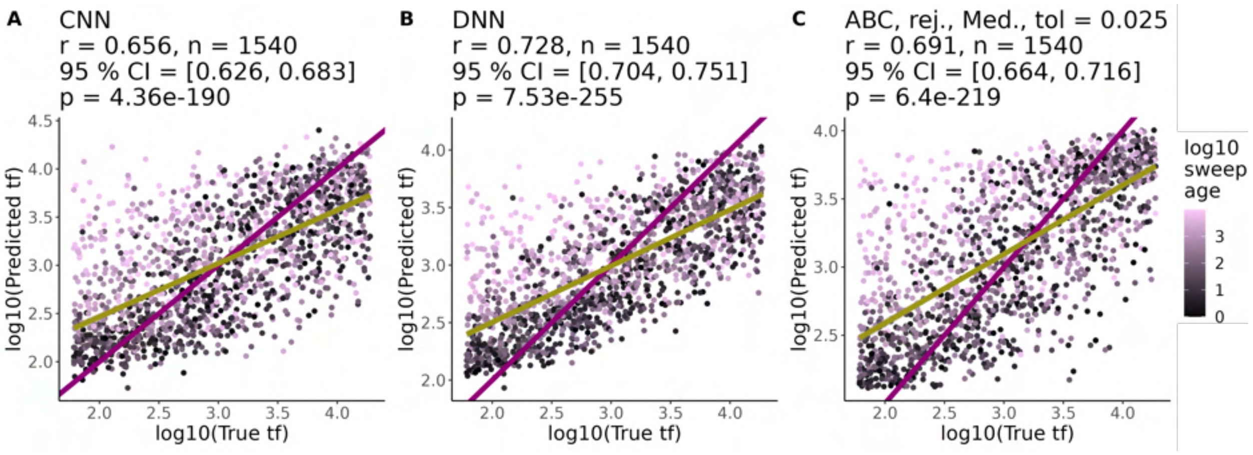

Figure S12: **Comparison of best CNN, DNN, and ABC models at predicting times to fixation for the cycle demography scenario.** Performance of best (A) CNN, (B) DNN, and (C) ABC models at predicting $t_f$ under the cycle demographic scenario. For each model, we list Pearson correlation coefficients between predicted and true values ($r$), the number of test set examples ($n$), the 95 % confidence intervals for $r$, and the p-value for testing if $r$ is different from 0 ($p$). The purple line is the 1-1 reference line where predictions and truth are equal. The yellow line is a least squares regression line. For the ABC model, the best performing model was a rejection regression, using the median of the posterior distribution as the point estimate, and a tolerance value of 0.025. All points are colored according to the age of the selective sweep in generations (log10 transformed).

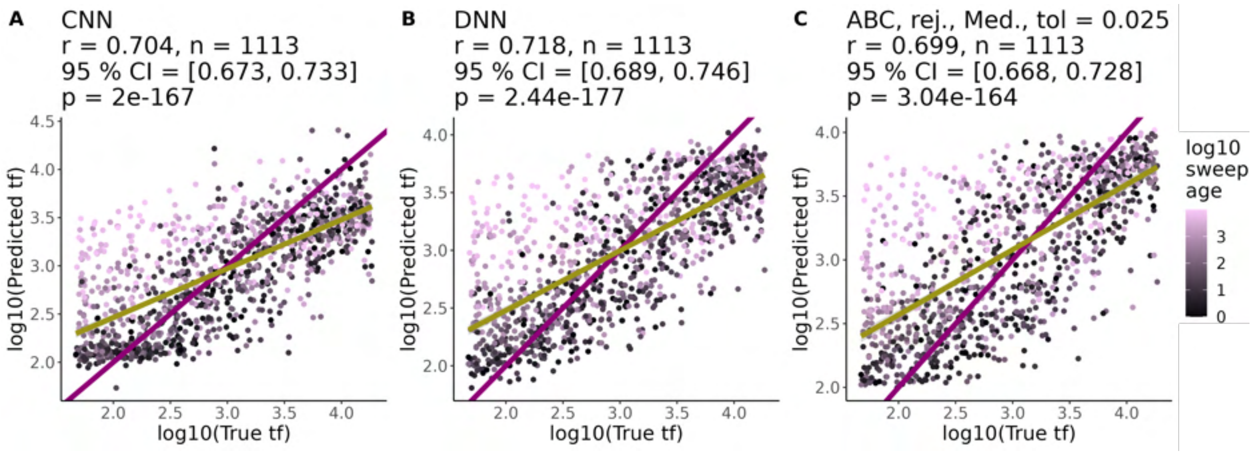

Figure S13: **Comparison of best models at predicting times to fixation for the chaos demography scenario.** Performance of best (A) CNN, (B) DNN, and (C) ABC models at predicting $t_f$ under the chaos demographic scenario. For each model, we list Pearson correlation coefficients between predicted and true values ($r$), the number of test set examples ($n$), the 95 % confidence intervals for $r$, and the p-value for testing if $r$ is different from 0 ($p$). The purple line is the 1-1 reference line where predictions and truth are equal. The yellow line is a least squares regression line. For the ABC model, the best performing model was a rejection (rej.) regression, using the median of the posterior distribution as the point estimate, and a tolerance value of 0.025. All points are colored according to the age of the selective sweep in generations (log10 transformed).

#### 2.2 Batch 2: Simple linear models for predicting $t_f$ from one summary statistic

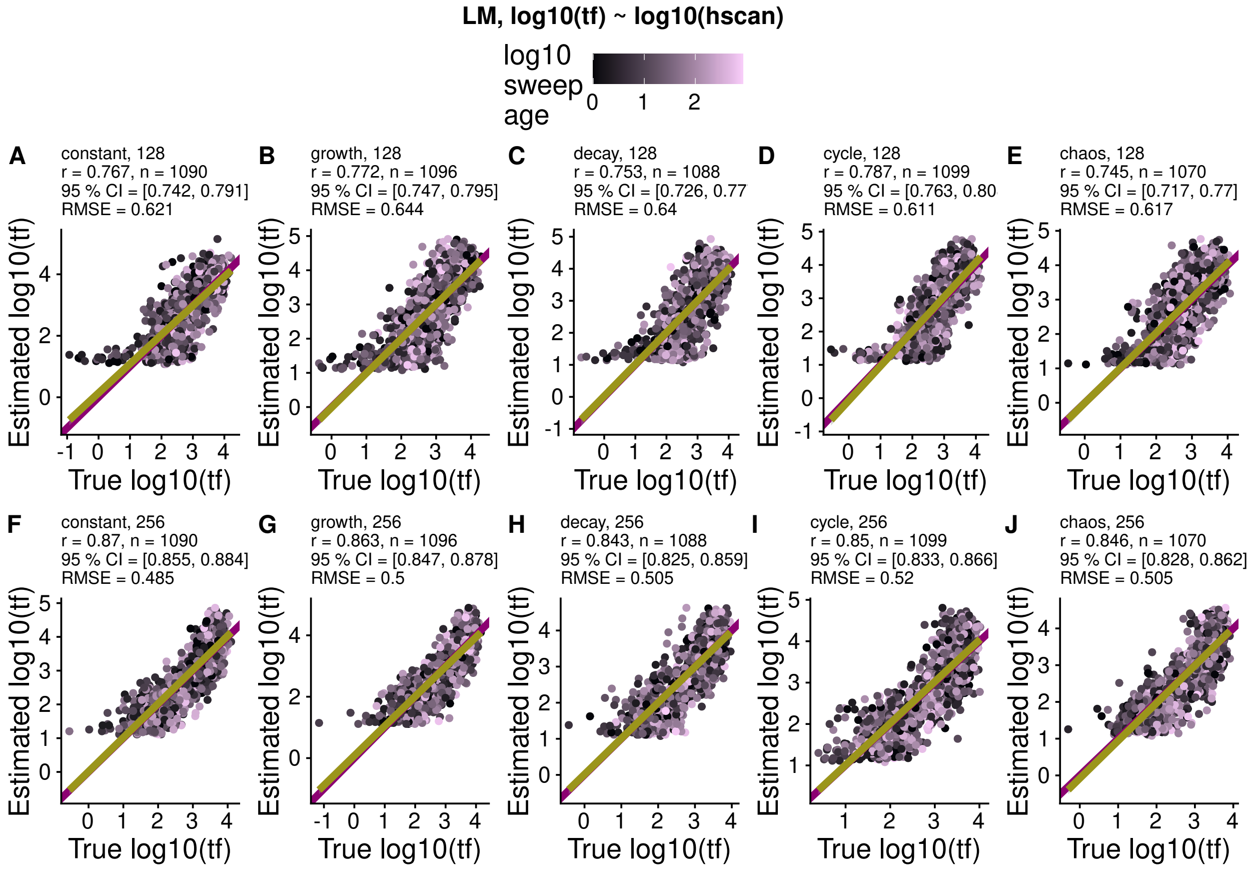

Figure S14: $t_f$ predictions from a simple linear model between $log10(t_f)$ and Messer's Hscan.

##### 2.3 Batch 2: Comparison of best CNN, DNN, and ABC models

| Outcome | Demography | m | n | $R^2$ | Adjusted $R^2$ |
|---|---|---|---|---|---|
| $log10(t_f + t_a)$ | chaos | 128 | 128 | 0.771 | 0.771 |
| $log10(t_f + t_a)$ | chaos | 256 | 128 | 0.795 | 0.795 |
| $log10(t_f + t_a)$ | constant | 128 | 128 | 0.813 | 0.813 |
| $log10(t_f + t_a)$ | constant | 256 | 128 | 0.828 | 0.827 |
| $log10(t_f + t_a)$ | cycle | 128 | 128 | 0.773 | 0.772 |
| $log10(t_f + t_a)$ | cycle | 256 | 128 | 0.79 | 0.79 |
| $log10(t_f + t_a)$ | decay | 128 | 128 | 0.78 | 0.779 |
| $log10(t_f + t_a)$ | decay | 256 | 128 | 0.801 | 0.801 |
| $log10(t_f + t_a)$ | growth | 128 | 128 | 0.839 | 0.839 |
| $log10(t_f + t_a)$ | growth | 256 | 128 | 0.849 | 0.849 |
| $log10(t_f)$ | chaos | 128 | 128 | 0.787 | 0.787 |
| $log10(t_f)$ | chaos | 256 | 128 | 0.817 | 0.816 |
| $log10(t_f)$ | constant | 128 | 128 | 0.832 | 0.831 |
| $log10(t_f)$ | constant | 256 | 128 | 0.856 | 0.855 |
| $log10(t_f)$ | cycle | 128 | 128 | 0.828 | 0.828 |
| $log10(t_f)$ | cycle | 256 | 128 | 0.851 | 0.85 |
| $log10(t_f)$ | decay | 128 | 128 | 0.8 | 0.8 |
| $log10(t_f)$ | decay | 256 | 128 | 0.828 | 0.828 |
| $log10(t_f)$ | growth | 128 | 128 | 0.861 | 0.86 |
| $log10(t_f)$ | growth | 256 | 128 | 0.879 | 0.879 |

Table S1: Total $R^2$ for predicting the outcomes $\log_{10}(t_f + t_a)$ and $\log_{10}(t_f)$ from summary statistics alone.

| **Demography** | **SNP window** | **ABC** | **DNN** | **CNN** | **CNN+DNN** |
|---|---|---|---|---|---|
| constant | 128 | 0.442 | 0.443 | 0.446 | 0.461 |
| growth | 128 | 0.459 | 0.443 | 0.570 | 0.453 |
| decay | 128 | 0.443 | 0.492 | 0.504 | 0.488 |
| cycle | 128 | 0.436 | 0.434 | 0.507 | 0.472 |
| chaos | 128 | 0.473 | 0.466 | 0.480 | 0.544 |
| constant | 256 | 0.423 | 0.422 | 0.448 | 0.472 |
| growth | 256 | 0.381 | 0.433 | 0.438 | 0.456 |
| decay | 256 | 0.424 | 0.441 | 0.498 | 0.468 |
| cycle | 256 | 0.424 | 0.486 | 0.470 | 0.517 |
| chaos | 256 | 0.456 | 0.434 | 0.457 | 0.496 |

Table S2: Root mean square error (RMSE) of $\log_{10}(t_f)$ predictions on the held-out test sets for the best-performing ABC, DNN, CNN, and CNN+DNN models in **Batch 2** simulations, for each demographic scenario and SNP window length. Values are those shown in Figure S15-S19 (128 SNP window) and Figures 4 and S20-S23 (256 SNP window). The horizontal rule separates the 128 SNP window scenarios (rows 2-6) from the 256 SNP window scenarios.

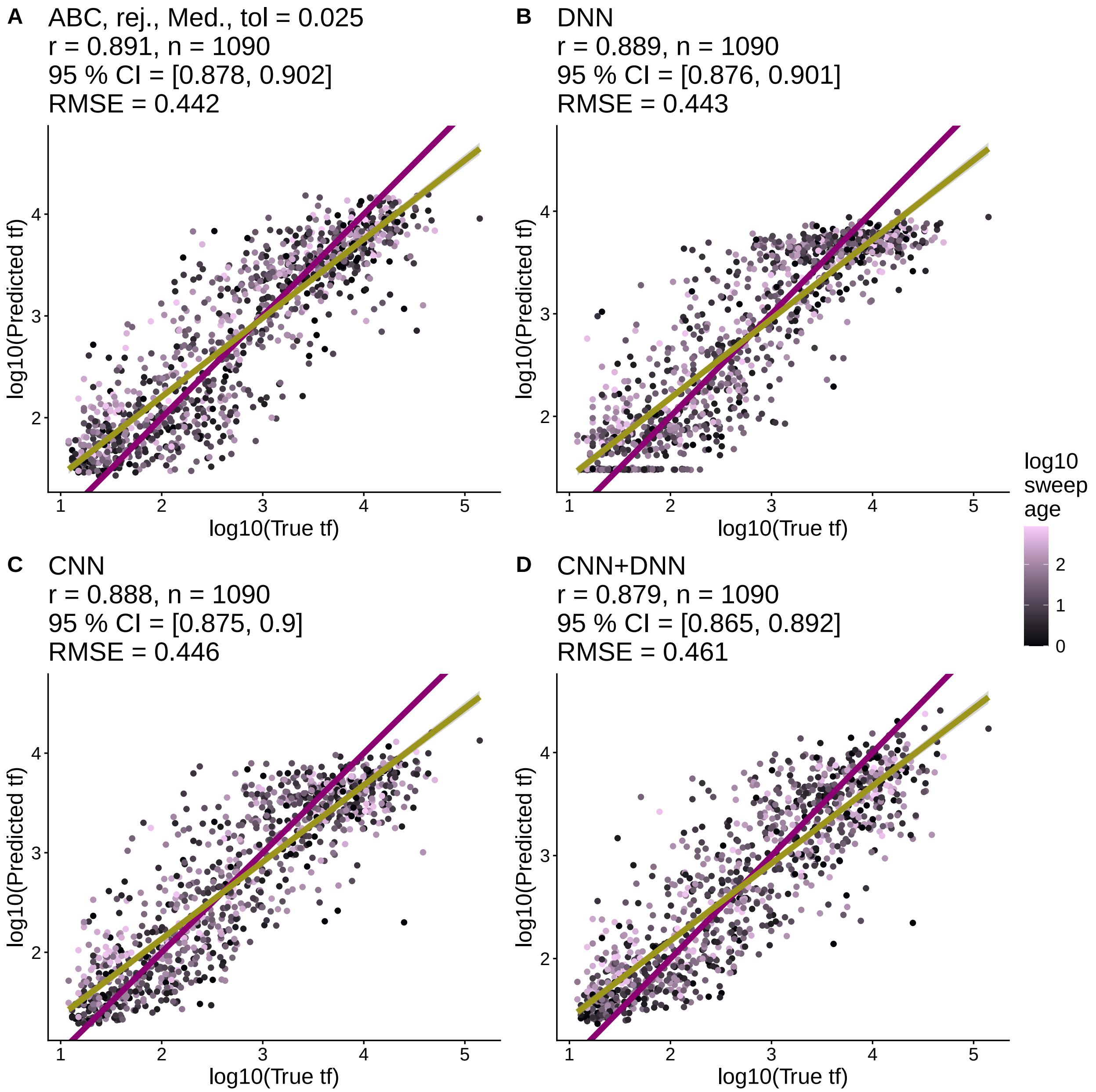

Figure S15: **Performance of CNN, DNN, and ABC models in Batch 2 simulations, 128 SNP window, constant demographic scenario.** Comparisons show true vs predicted $t_f$ for best-performing (A) ABC, (B) DNN, (C) CNN, and (D) CNN+DNN models. The best performing ABC model used the rejection method (rej.), a median posterior estimate (Med.) and a tolerance value of 0.025 (tol). For each model, we list Pearson correlation coefficients between predicted and true values ($r$), the number of test set examples ($n$), the 95 % confidence intervals for $r$, the p-value for testing if $r$ is different from 0 ($p$), and the root mean square error of the predictions from ground truth (RMSE). The purple line is the 1-1 reference line where predictions and truth are equal. The yellow line is a least squares regression line. All points are colored according to the $t_a$ of the sweep in generations (log10 transformed).

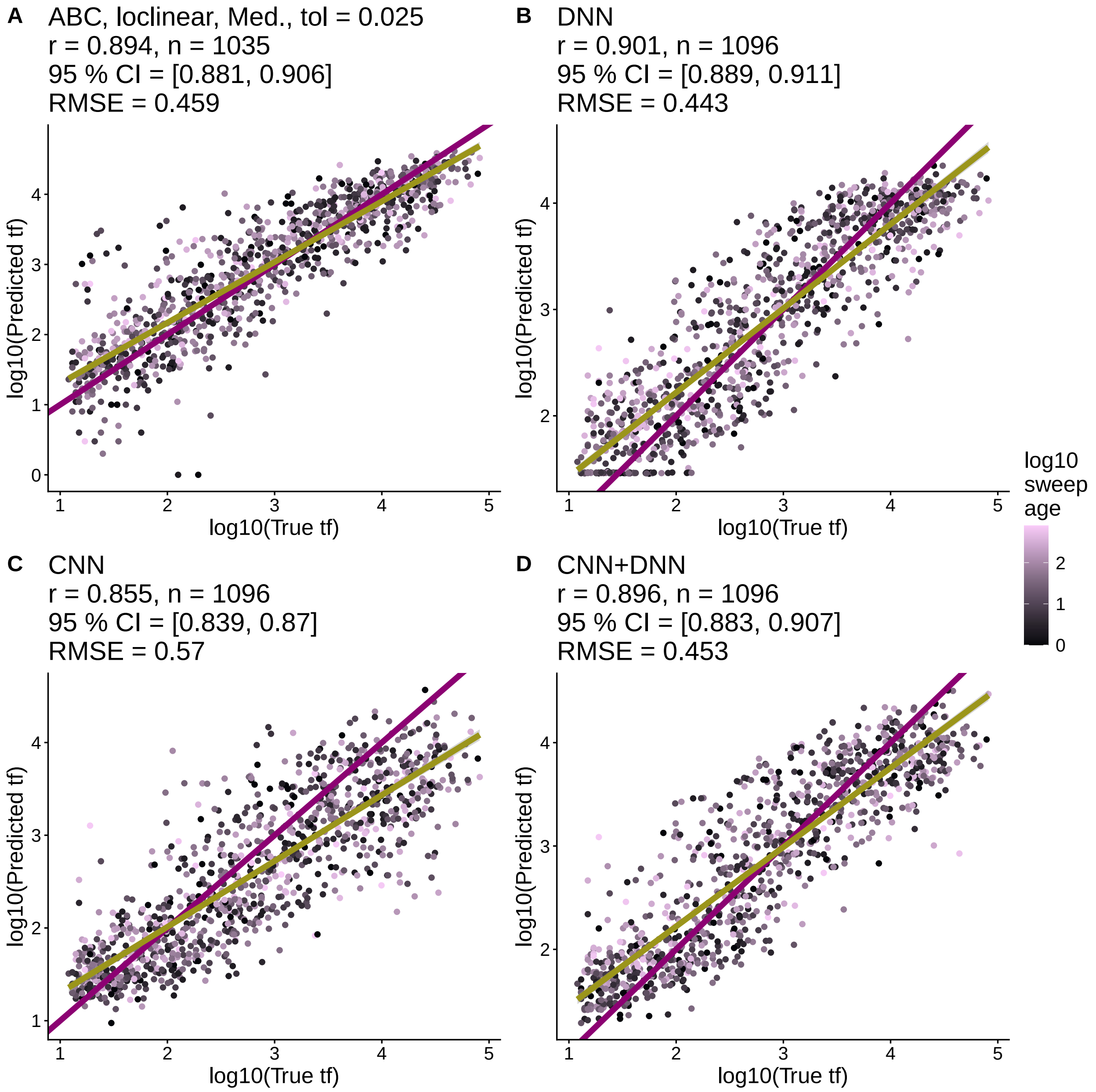

Figure S16: **Performance of CNN, DNN, and ABC models in Batch 2 simulations, 128 SNP window, growth demographic scenario.** The best performing ABC model used the rejection method (rej.), a median posterior estimate (Med.) and a tolerance value of 0.025 (tol). For each model, we list Pearson correlation coefficients between predicted and true values ($r$), the number of test set examples ($n$), the 95 % confidence intervals for $r$, the p-value for testing if $r$ is different from 0 ($p$), and the root mean square error of the predictions from ground truth (RMSE). The purple line is the 1-1 reference line where predictions and truth are equal. The yellow line is a least squares regression line. All points are colored according to the $t_a$ of the sweep in generations (log10 transformed).

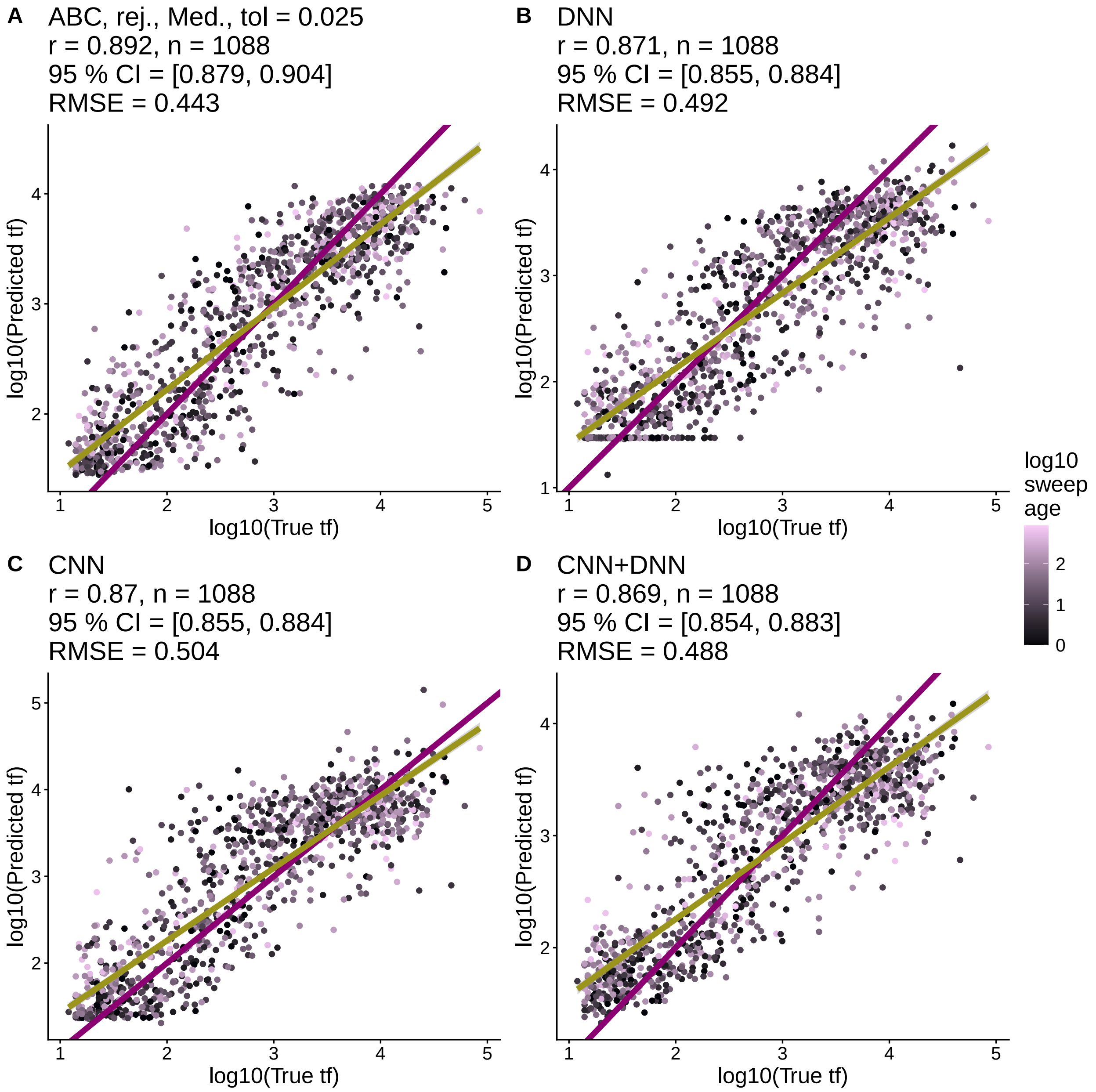

Figure S17: **Performance of CNN, DNN, and ABC models in Batch 2 simulations, 128 SNP window, decay demographic scenario.** The best performing ABC model used the rejection method (rej.), a median posterior estimate (Med.) and a tolerance value of 0.025 (tol). For each model, we list Pearson correlation coefficients between predicted and true values ($r$), the number of test set examples ($n$), the 95 % confidence intervals for $r$, the p-value for testing if $r$ is different from 0 ($p$), and the root mean square error of the predictions from ground truth (RMSE). The purple line is the 1-1 reference line where predictions and truth are equal. The yellow line is a least squares regression line. All points are colored according to the $t_a$ of the sweep in generations (log10 transformed).

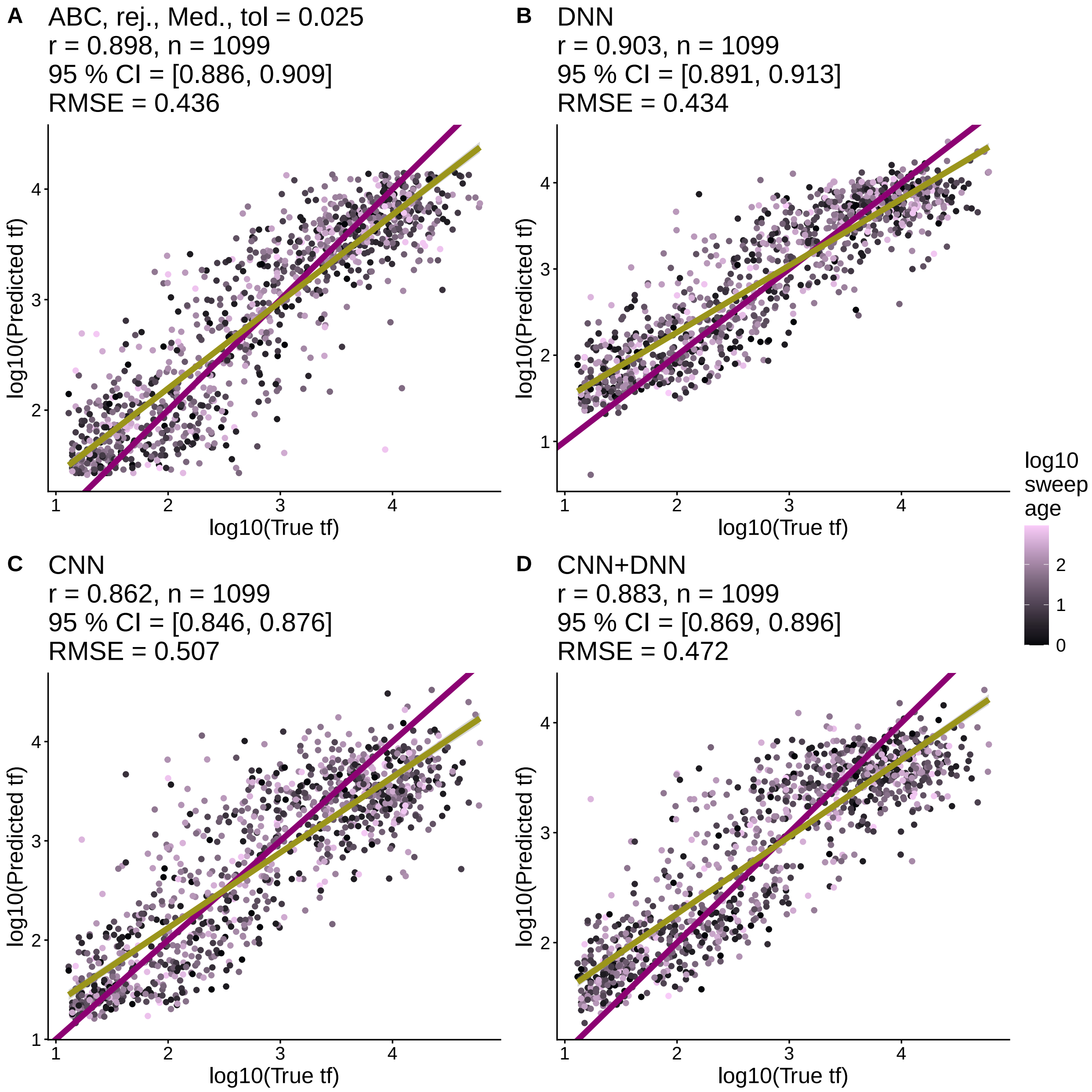

Figure S18: **Performance of CNN, DNN, and ABC models in Batch 2 simulations, 128 SNP window, cycle demographic scenario.** The best performing ABC model used the rejection method (rej.), a median posterior estimate (Med.) and a tolerance value of 0.025 (tol). For each model, we list Pearson correlation coefficients between predicted and true values ($r$), the number of test set examples ($n$), the 95 % confidence intervals for $r$, the p-value for testing if $r$ is different from 0 ($p$), and the root mean square error of the predictions from ground truth (RMSE). The purple line is the 1-1 reference line where predictions and truth are equal. The yellow line is a least squares regression line. All points are colored according to the $t_a$ of the sweep in generations (log10 transformed).

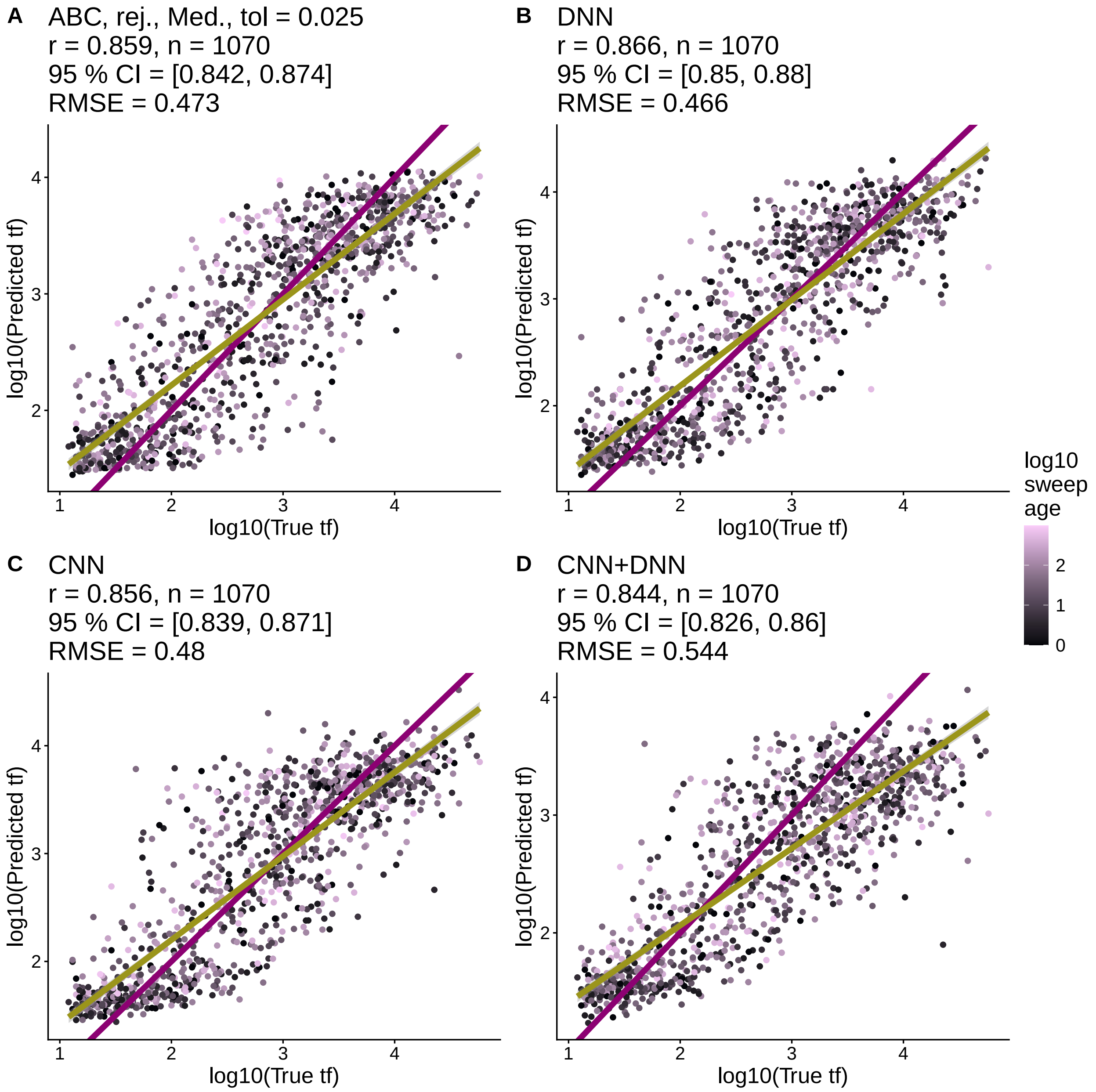

Figure S19: **Performance of CNN, DNN, and ABC models in Batch 2 simulations, 128 SNP window, chaos demographic scenario.** The best performing ABC model used the rejection method (rej.), a median posterior estimate (Med.) and a tolerance value of 0.025 (tol). For each model, we list Pearson correlation coefficients between predicted and true values ($r$), the number of test set examples ($n$), the 95 % confidence intervals for $r$, the p-value for testing if $r$ is different from 0 ($p$), and the root mean square error of the predictions from ground truth (RMSE). The purple line is the 1-1 reference line where predictions and truth are equal. The yellow line is a least squares regression line. All points are colored according to the $t_a$ of the sweep in generations (log10 transformed).

#### 2.4 Batch 2: Comparison of best CNN, DNN, and ABC models, larger window size

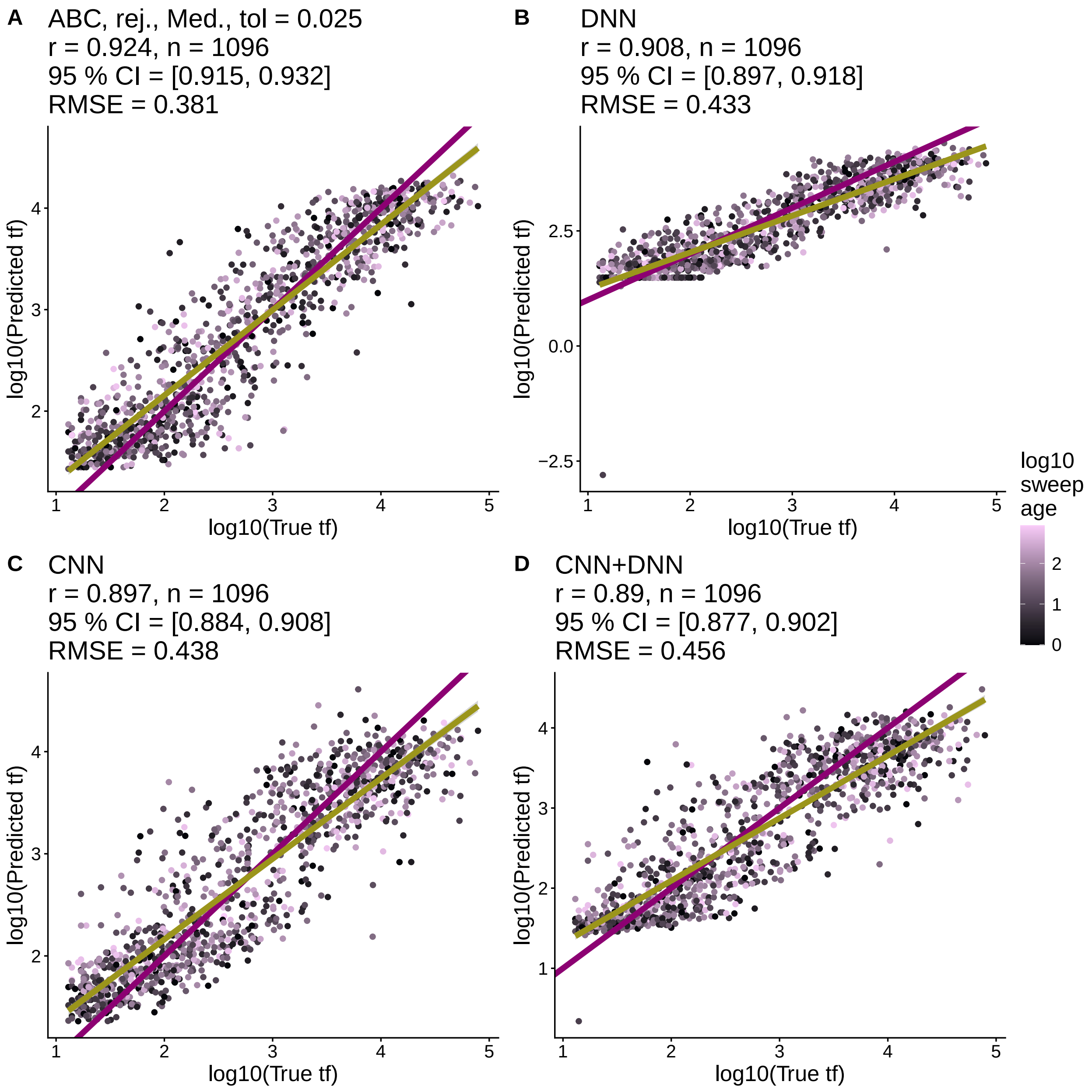

Figure S20: **Performance of CNN, DNN, and ABC models in Batch 2 simulations, 256 SNP window, growth demographic scenario.** The best performing ABC model used the rejection method (rej.), a median posterior estimate (Med.) and a tolerance value of 0.025 (tol). For each model, we list Pearson correlation coefficients between predicted and true values ($r$), the number of test set examples ($n$), the 95 % confidence intervals for $r$, the p-value for testing if $r$ is different from 0 ($p$), and the root mean square error of the predictions from ground truth (RMSE). The purple line is the 1-1 reference line where predictions and truth are equal. The yellow line is a least squares regression line. All points are colored according to the $t_a$ of the sweep in generations (log10 transformed).

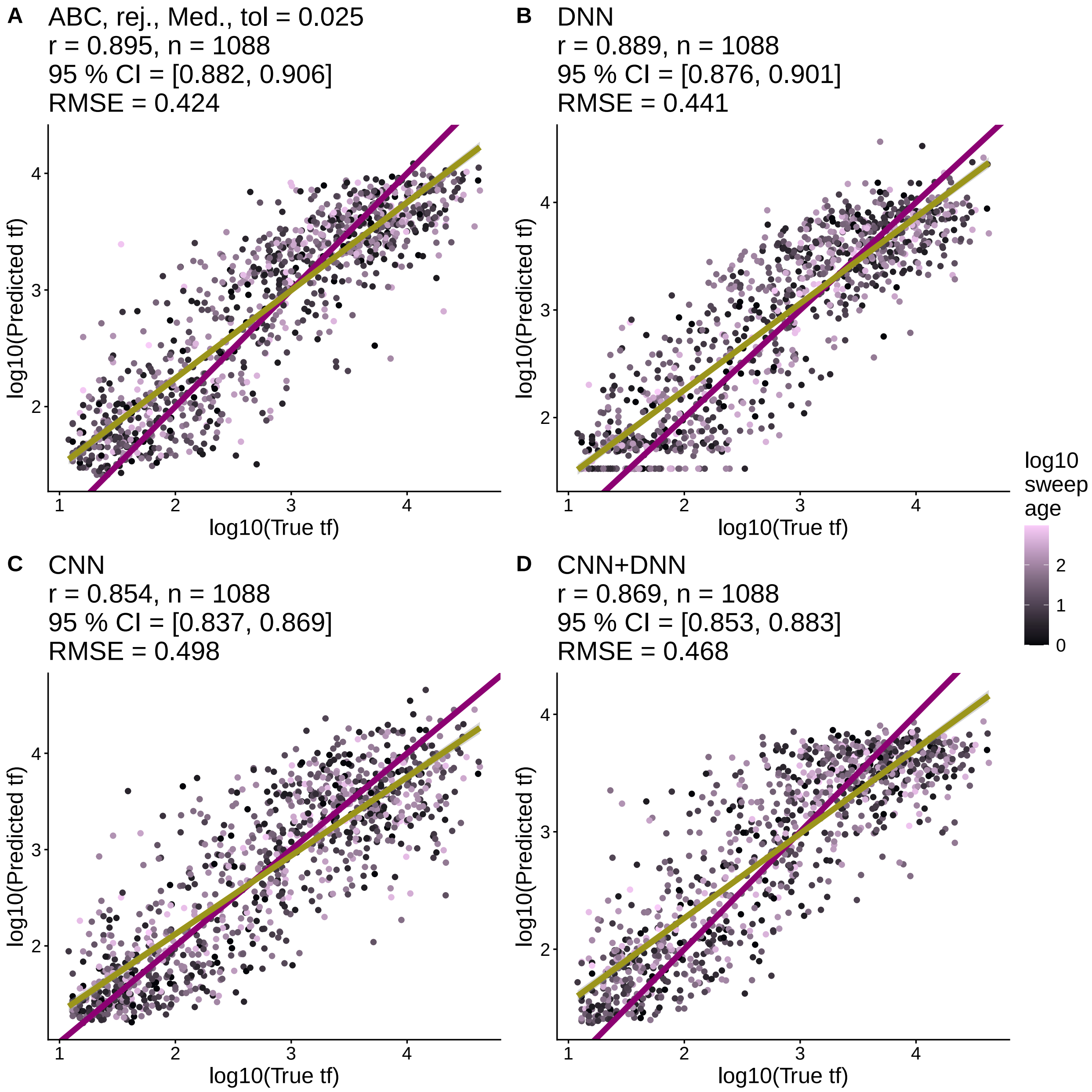

Figure S21: **Performance of CNN, DNN, and ABC models in Batch 2 simulations, 256 SNP window, decay demographic scenario.** The best performing ABC model used the rejection method (rej.), a median posterior estimate (Med.) and a tolerance value of 0.025 (tol). For each model, we list Pearson correlation coefficients between predicted and true values ($r$), the number of test set examples ($n$), the 95 % confidence intervals for $r$, the p-value for testing if $r$ is different from 0 ($p$), and the root mean square error of the predictions from ground truth (RMSE). The purple line is the 1-1 reference line where predictions and truth are equal. The yellow line is a least squares regression line. All points are colored according to the $t_a$ of the sweep in generations (log10 transformed).

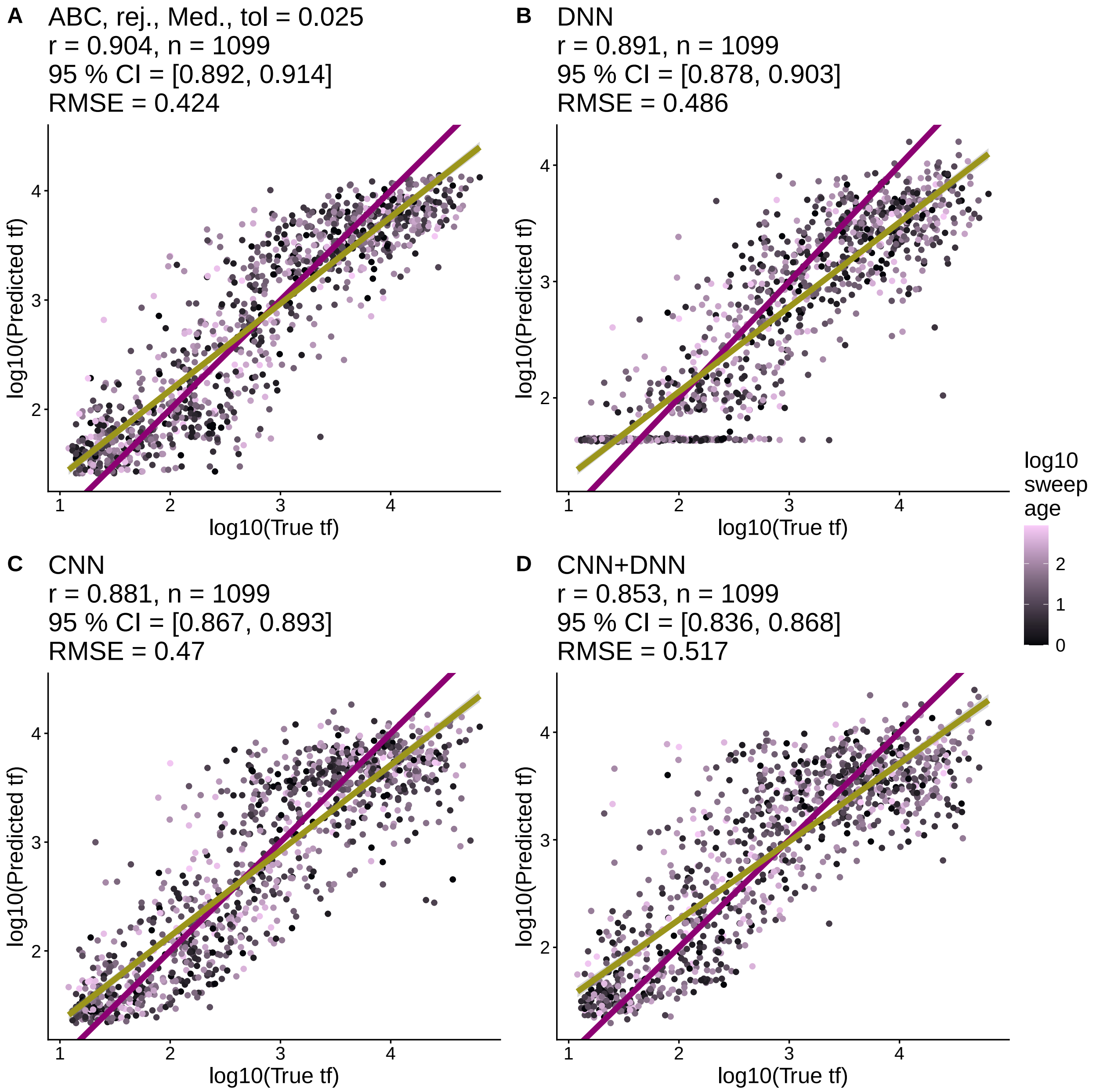

Figure S22: **Performance of CNN, DNN, and ABC models in Batch 2 simulations, 256 SNP window, cycle demographic scenario.** The best performing ABC model used the rejection method (rej.), a median posterior estimate (Med.) and a tolerance value of 0.025 (tol). For each model, we list Pearson correlation coefficients between predicted and true values ($r$), the number of test set examples ($n$), the 95 % confidence intervals for $r$, the p-value for testing if $r$ is different from 0 ($p$), and the root mean square error of the predictions from ground truth (RMSE). The purple line is the 1-1 reference line where predictions and truth are equal. The yellow line is a least squares regression line. All points are colored according to the $t_a$ of the sweep in generations (log10 transformed).

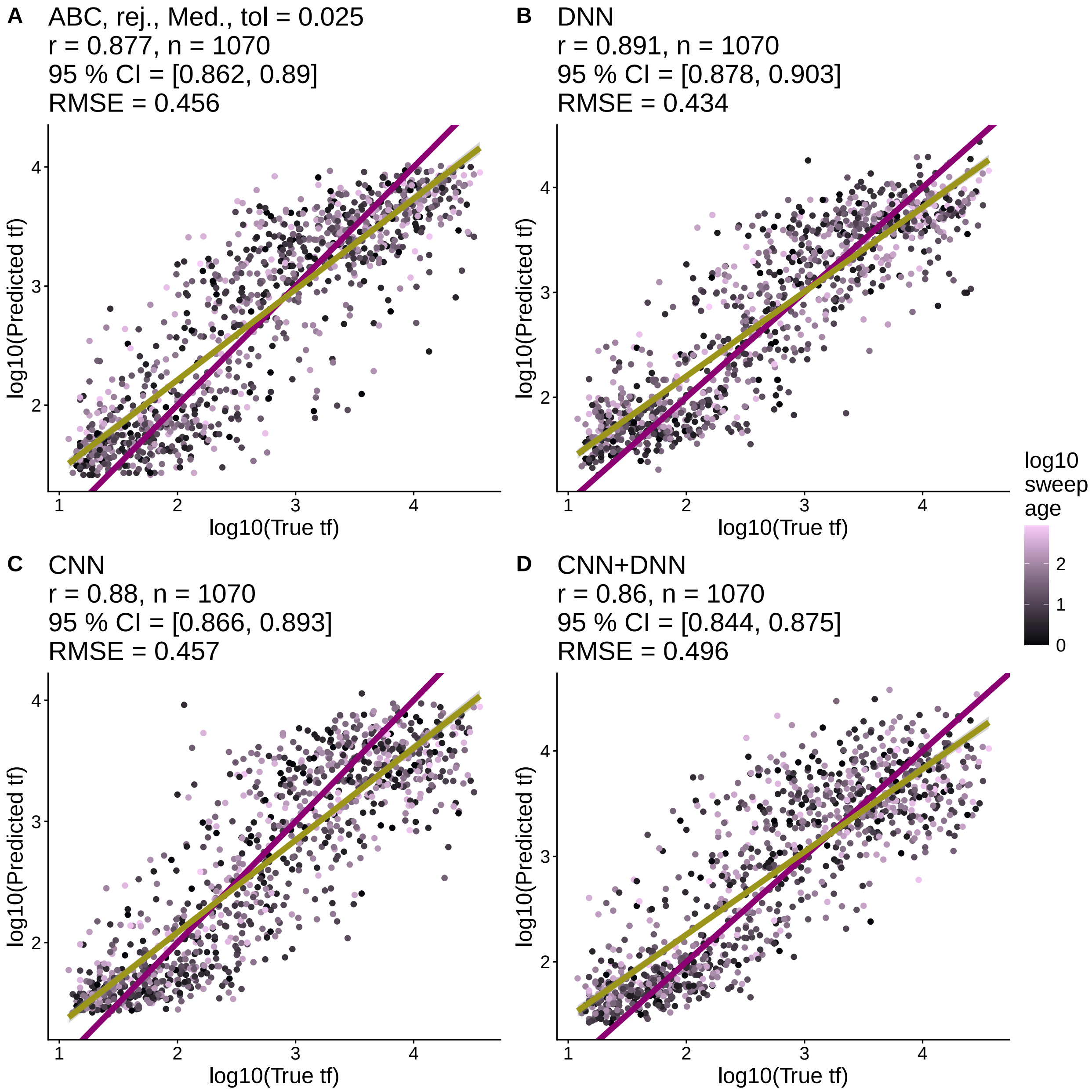

Figure S23: **Performance of CNN, DNN, and ABC models in Batch 2 simulations, 256 SNP window, chaos demographic scenario.** The best performing ABC model used the rejection method (rej.), a median posterior estimate (Med.) and a tolerance value of 0.025 (tol). For each model, we list Pearson correlation coefficients between predicted and true values ($r$), the number of test set examples ($n$), the 95 % confidence intervals for $r$, the p-value for testing if $r$ is different from 0 ($p$), and the root mean square error of the predictions from ground truth (RMSE). The purple line is the 1-1 reference line where predictions and truth are equal. The yellow line is a least squares regression line. All points are colored according to the $t_a$ of the sweep in generations (log10 transformed).

#### 2.5 Batch 2: Prediction error of best models

##### 2.5.1 128 SNP window

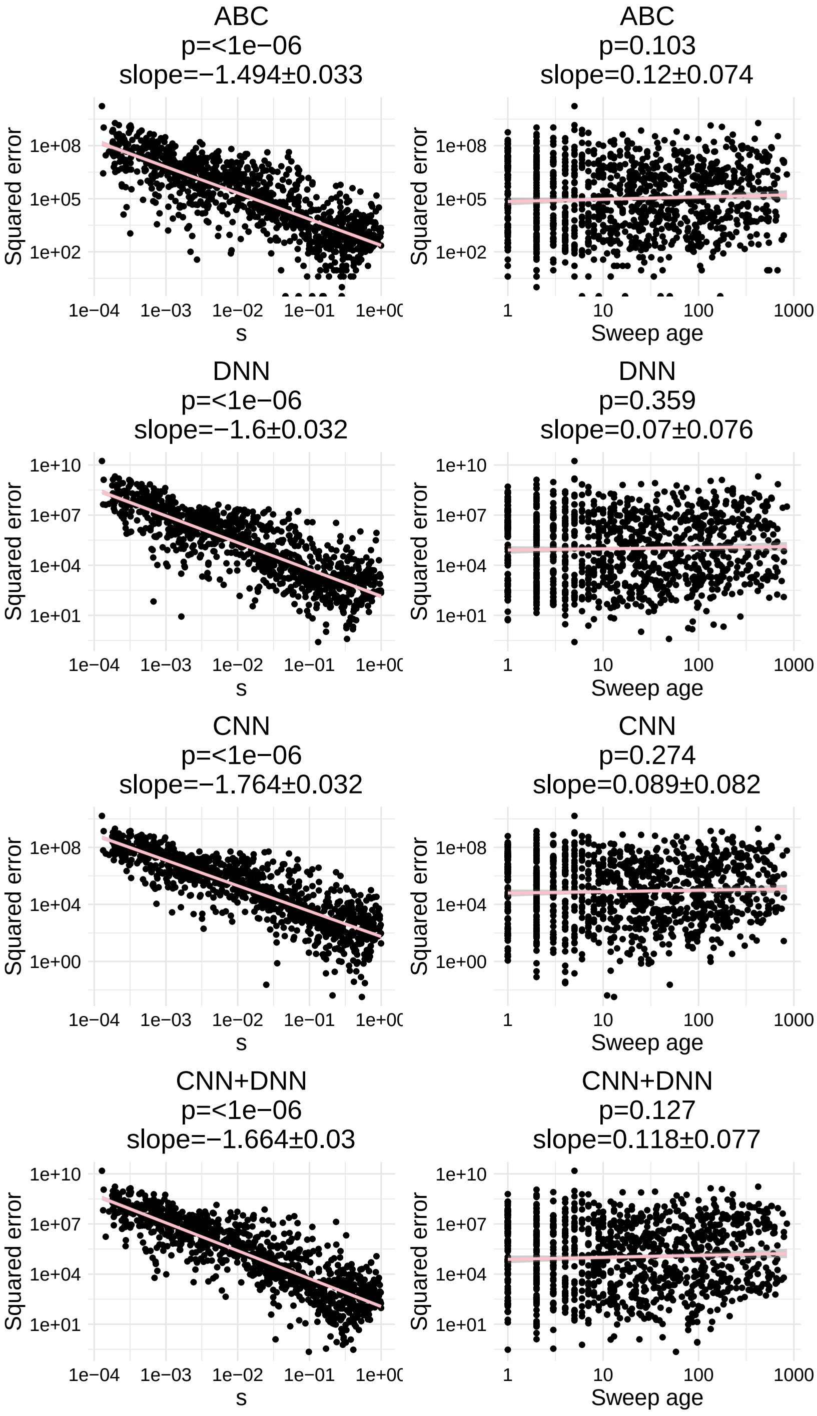

Figure S24: For the **constant** demographic scenario, a linear model analyzing the relationship between prediction error and key sweep parameters, $s$, which influences $t_f$, and sweep age $t_a$, which influences the degradation of the sweep signal.

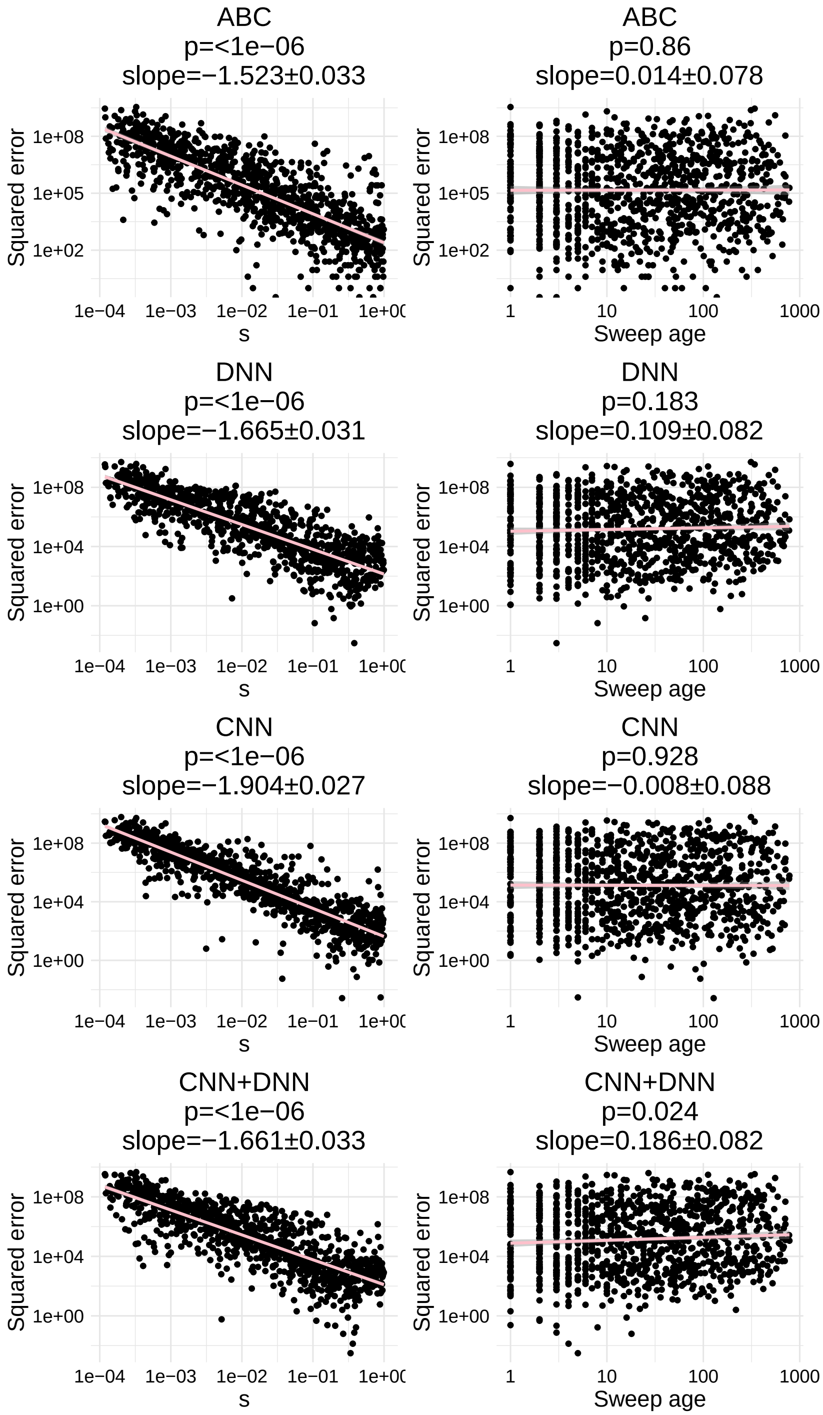

Figure S25: For the **growth** demographic scenario, a linear model analyzing the relationship between prediction error and key sweep parameters, $s$, which influences $t_f$, and sweep age $t_a$, which influences the degradation of the sweep signal.

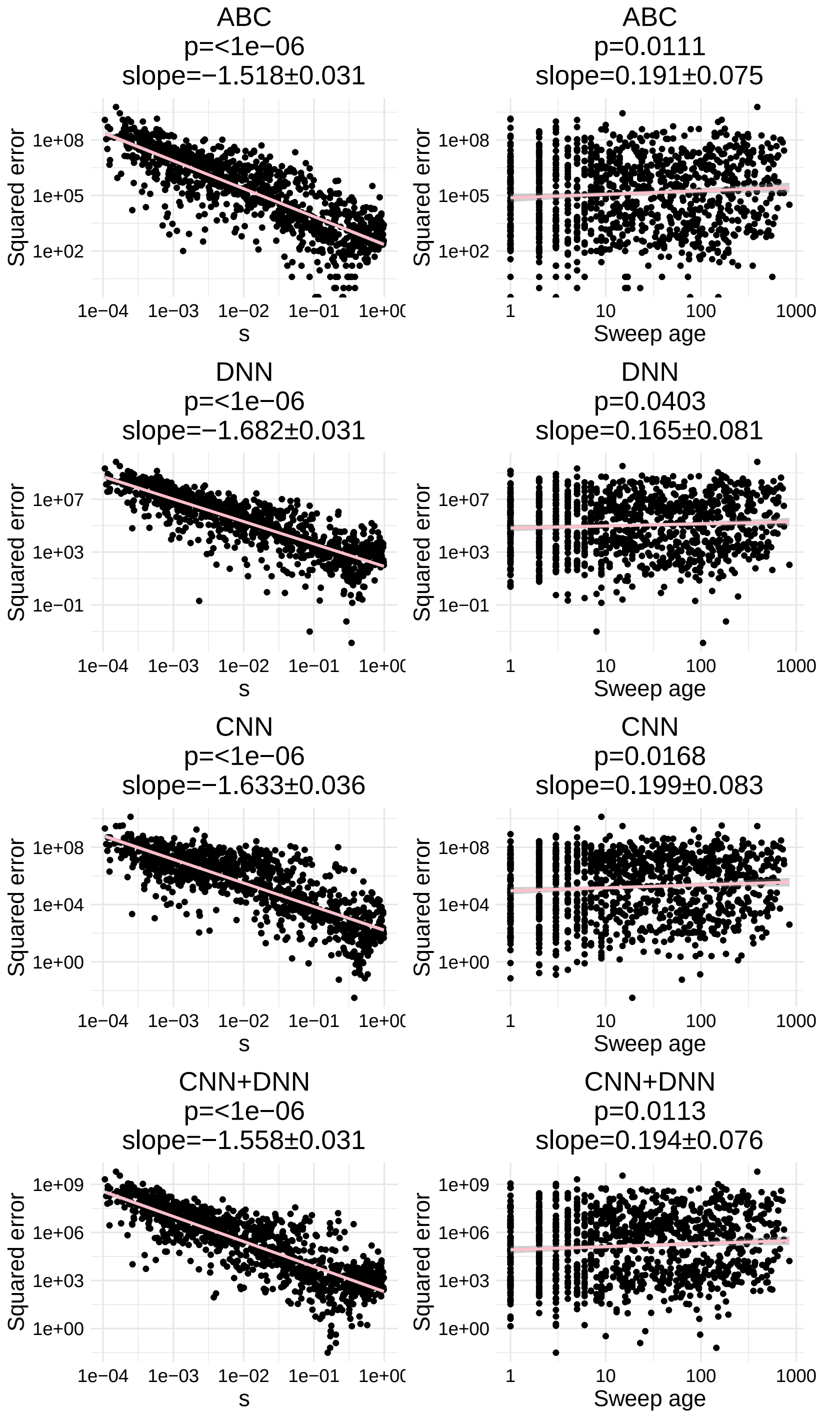

Figure S26: For the **decay** demographic scenario, a linear model analyzing the relationship between prediction error and key sweep parameters, $s$, which influences $t_f$, and sweep age $t_a$, which influences the degradation of the sweep signal.

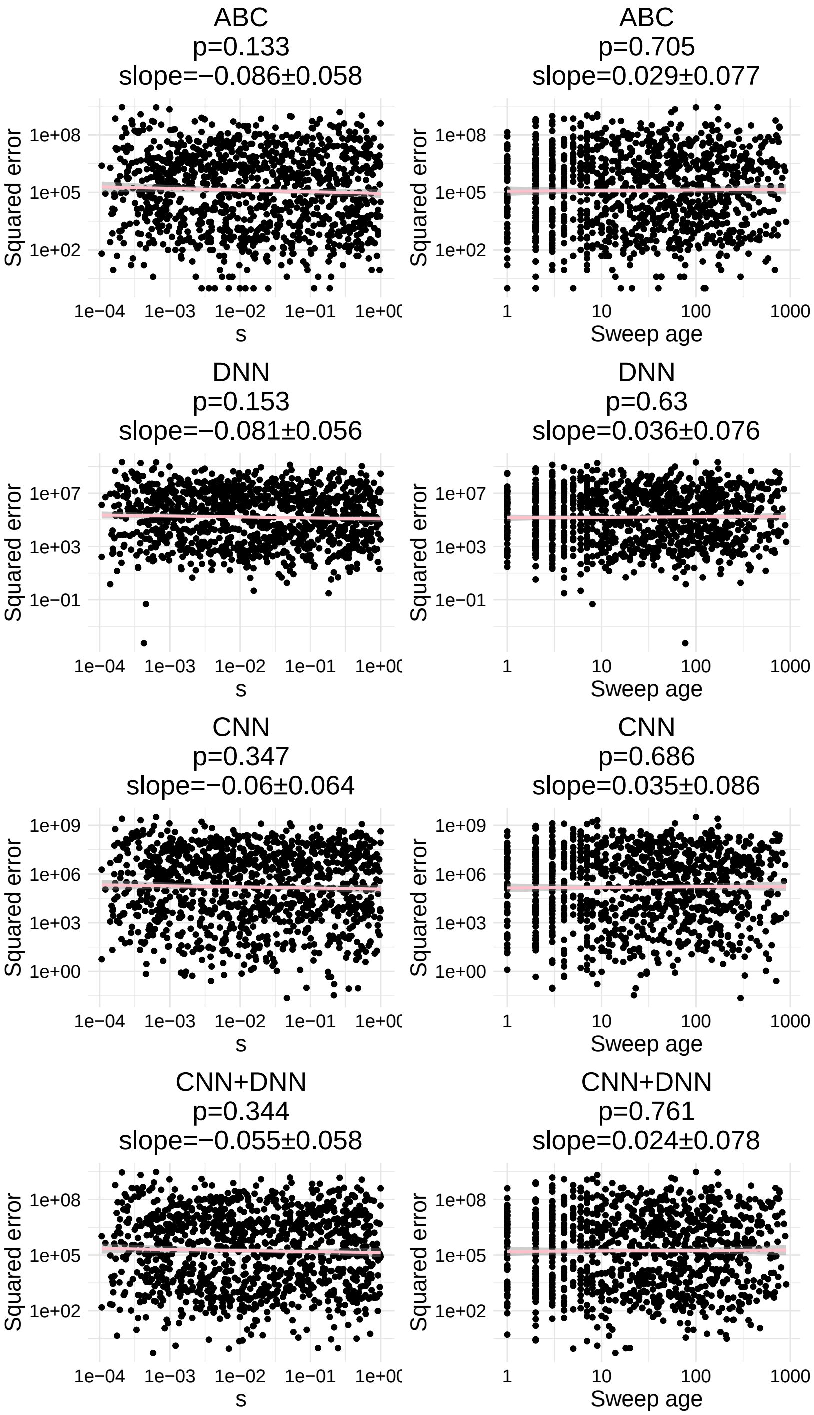

Figure S27: For the **cycle** demographic scenario, a linear model analyzing the relationship between prediction error and key sweep parameters, $s$, which influences $t_f$, and sweep age $t_a$, which influences the degradation of the sweep signal.

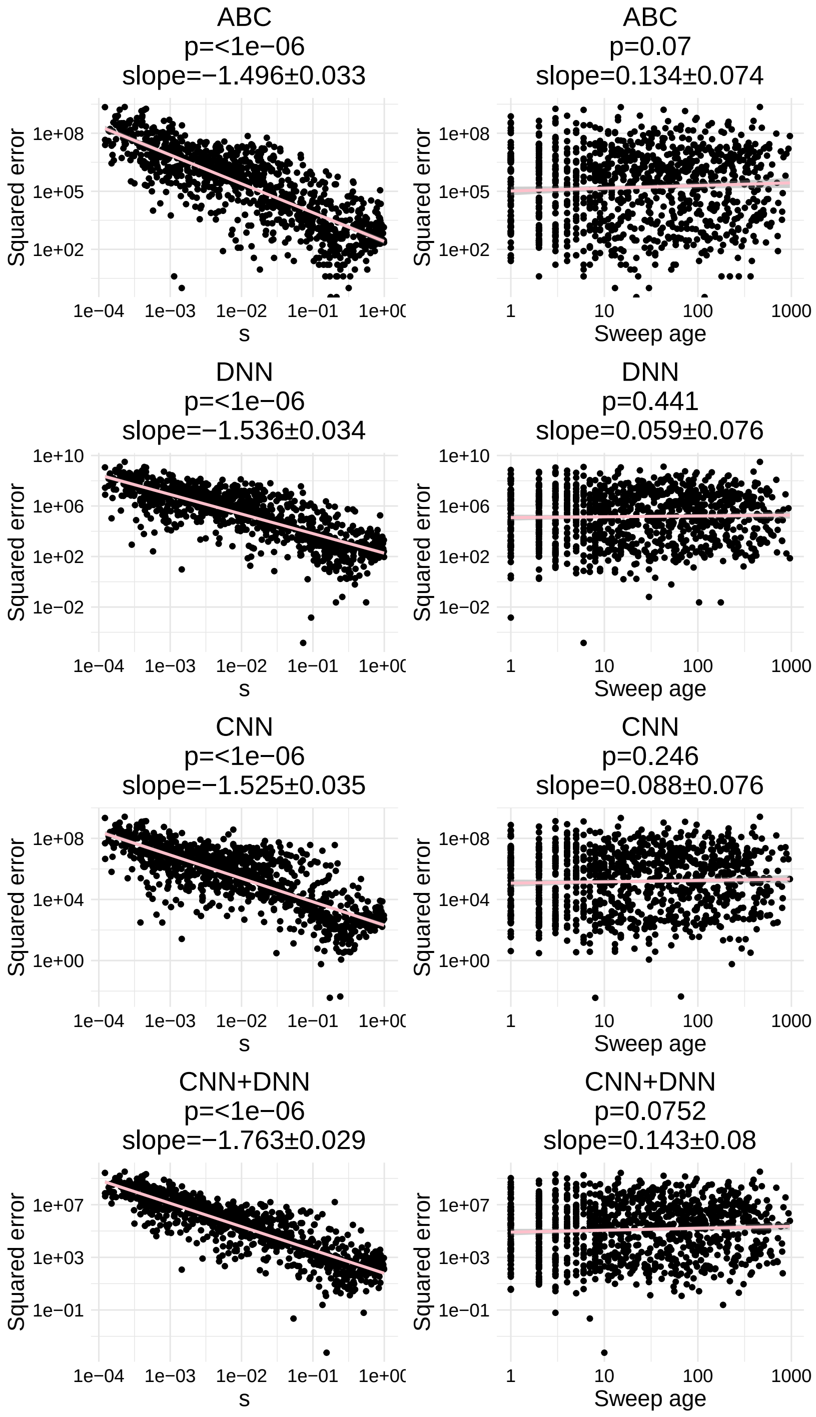

Figure S28: For the **chaos** demographic scenario, a linear model analyzing the relationship between prediction error and key sweep parameters, $s$, which influences $t_f$, and sweep age $t_a$, which influences the degradation of the sweep signal.

##### 2.5.2 256 SNP window

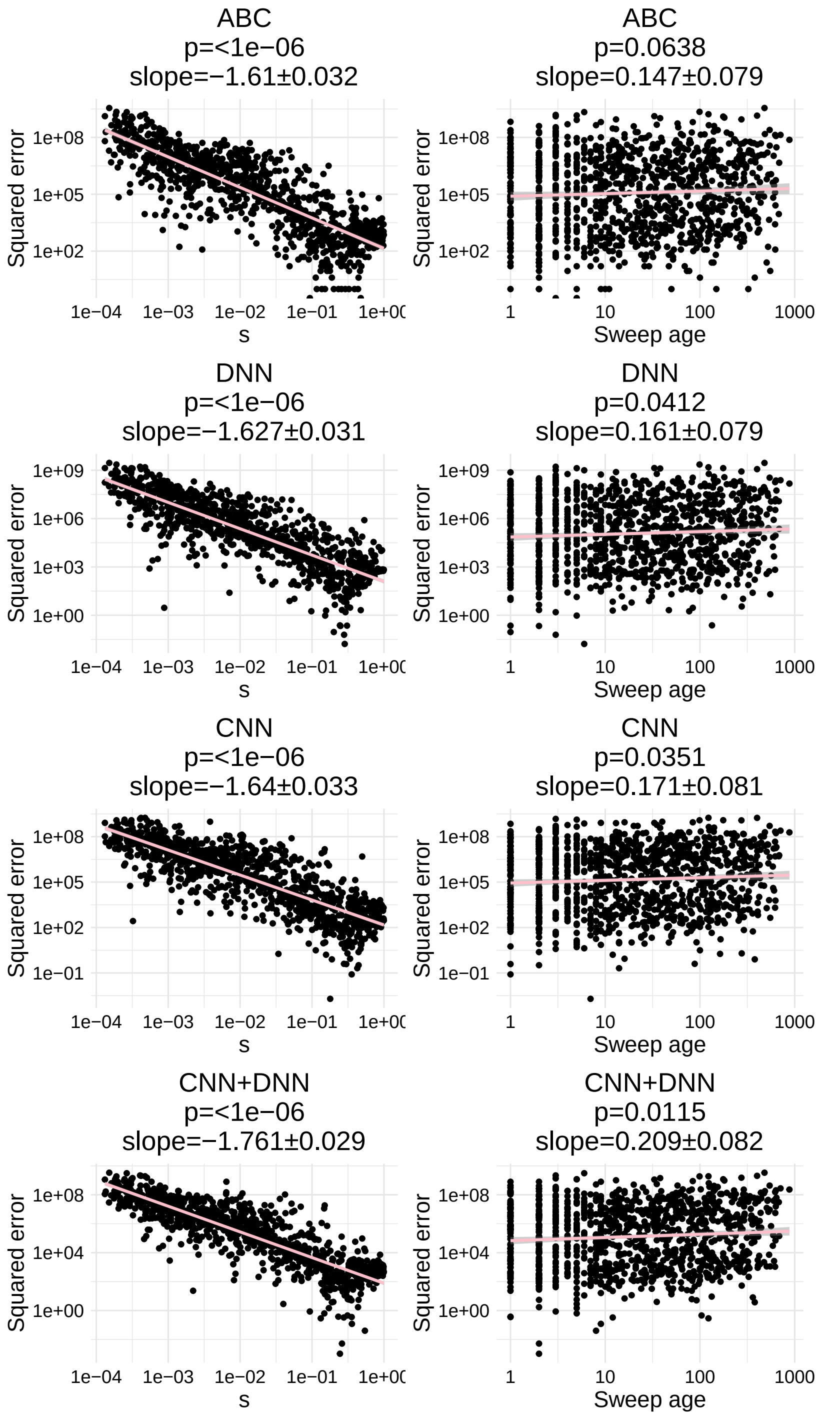

Figure S29: For the **constant** demographic scenario, a linear model analyzing the relationship between prediction error and key sweep parameters, $s$, which influences $t_f$, and sweep age $t_a$, which influences the degradation of the sweep signal.

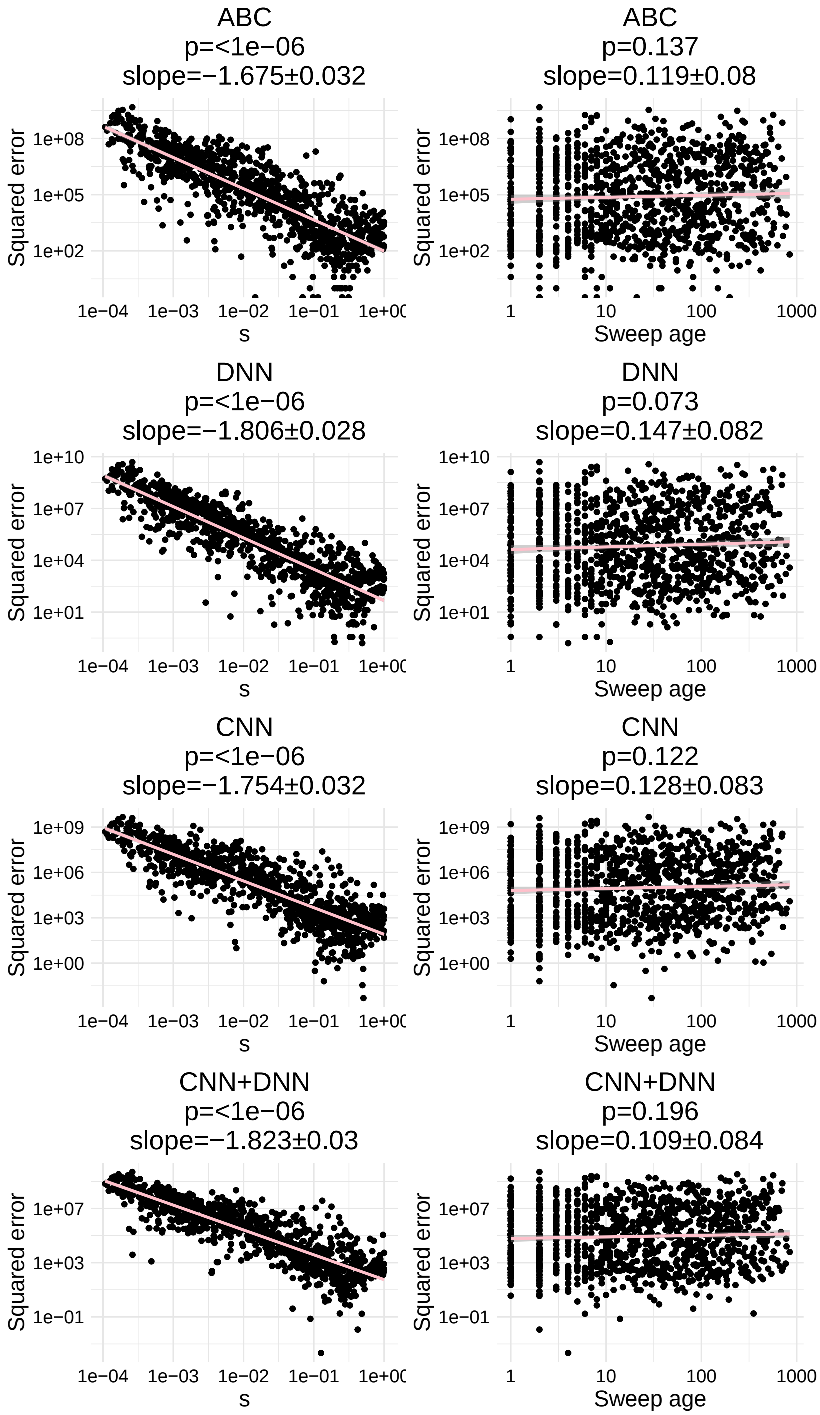

Figure S30: For the **growth** demographic scenario, a linear model analyzing the relationship between prediction error and key sweep parameters, $s$, which influences $t_f$, and sweep age $t_a$, which influences the degradation of the sweep signal.

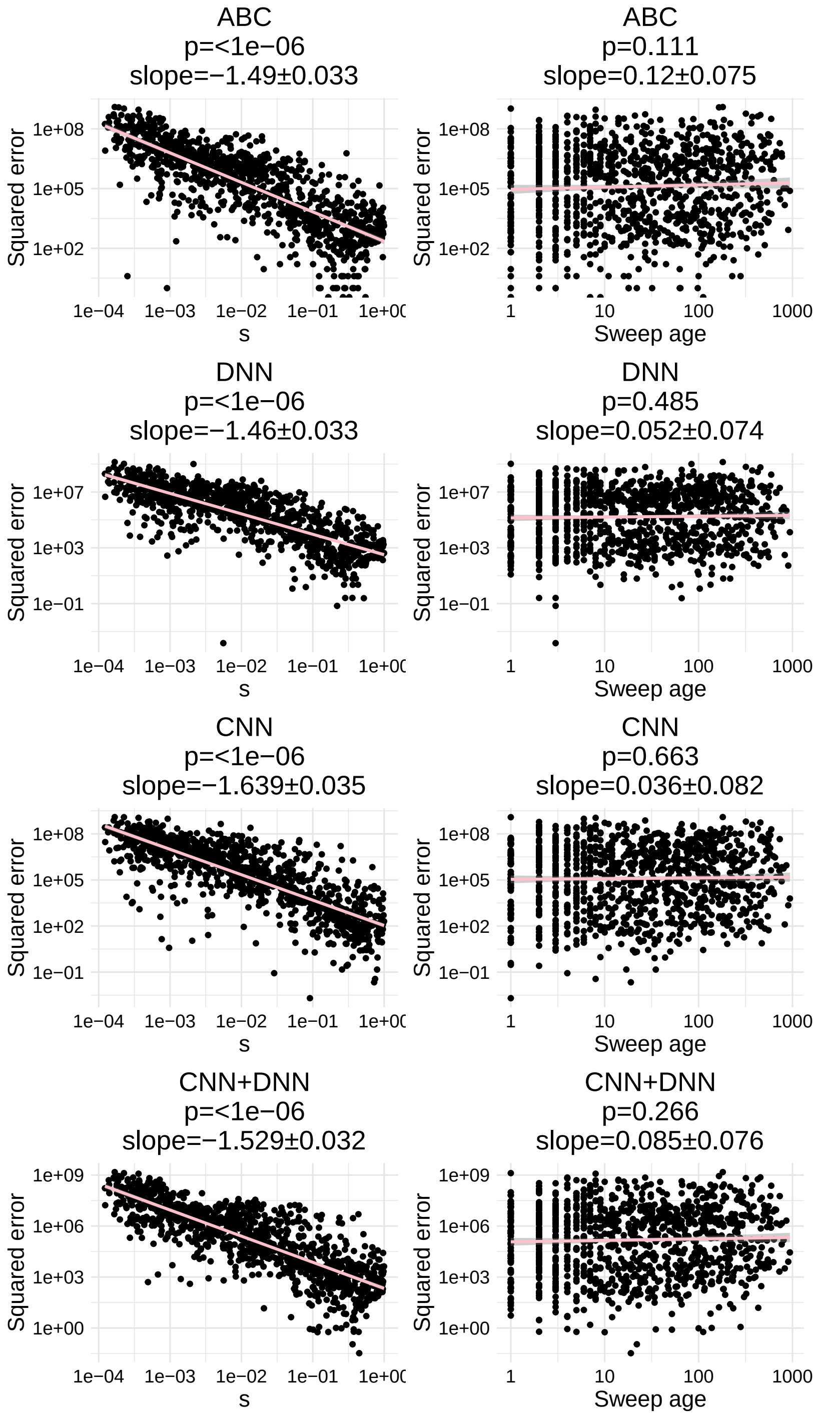

Figure S31: For the **decay** demographic scenario, a linear model analyzing the relationship between prediction error and key sweep parameters, $s$, which influences $t_f$, and sweep age $t_a$, which influences the degradation of the sweep signal.

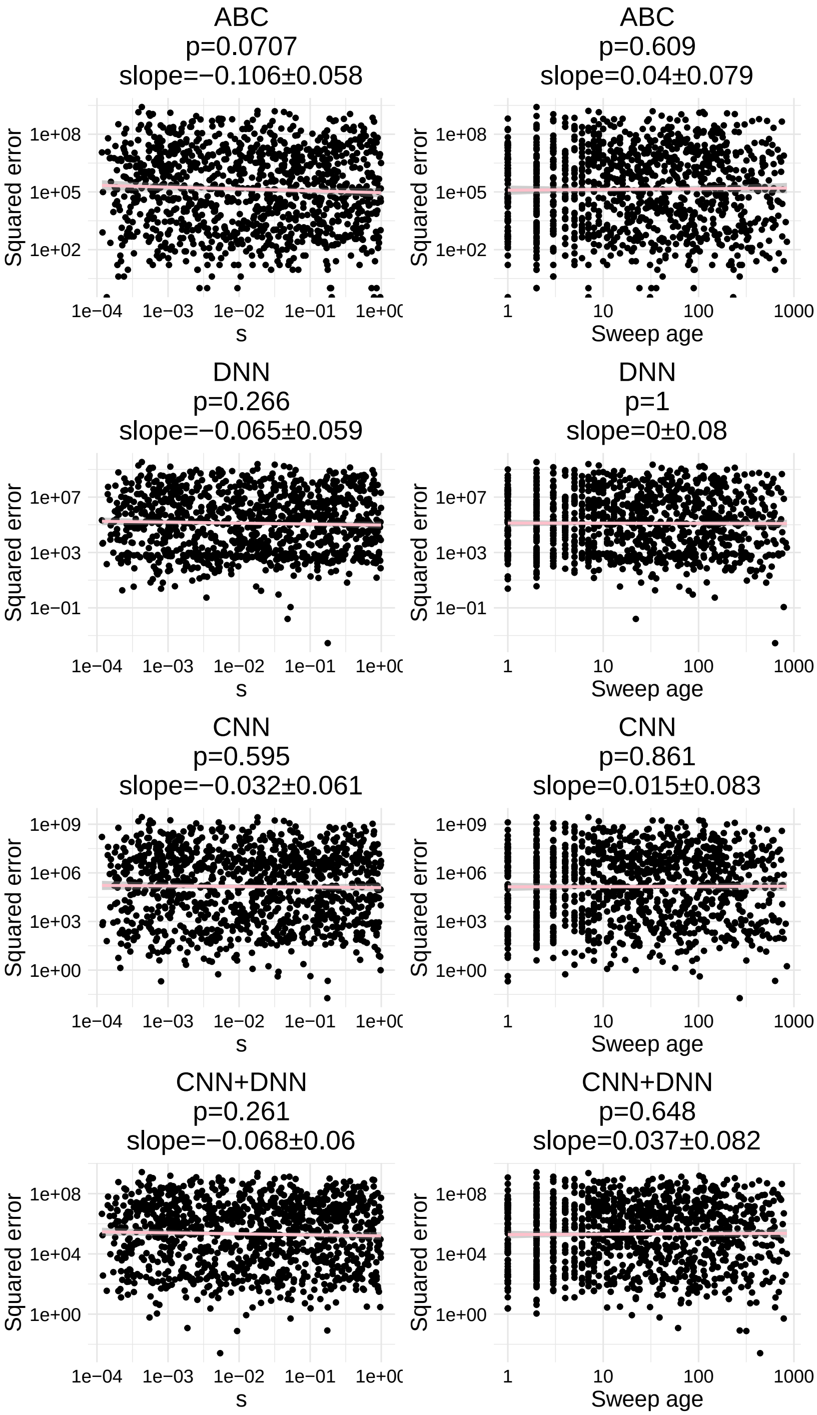

Figure S32: For the **cycle** demographic scenario, a linear model analyzing the relationship between prediction error and key sweep parameters, $s$, which influences $t_f$, and sweep age $t_a$, which influences the degradation of the sweep signal.

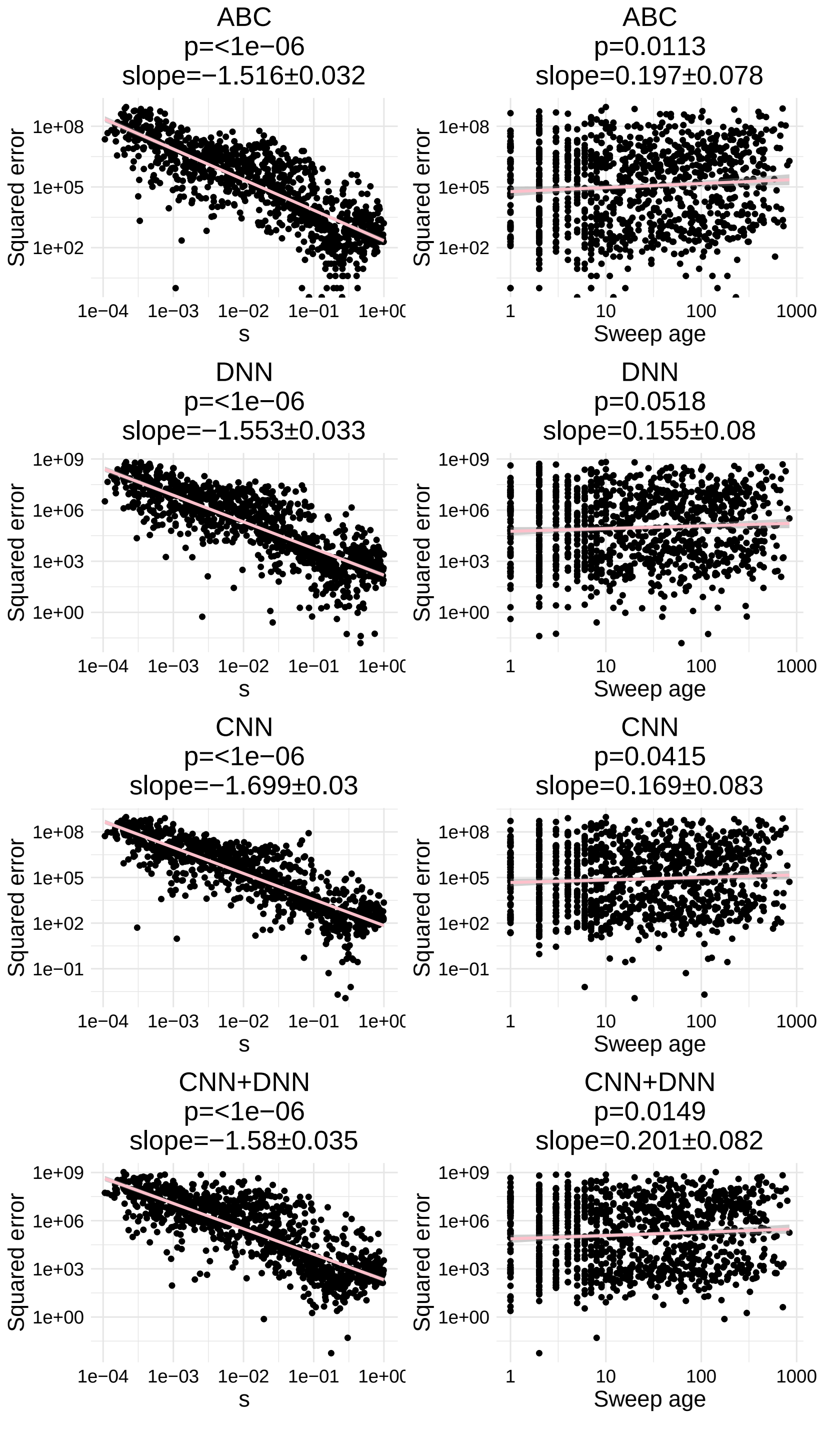

Figure S33: For the **chaos** demographic scenario, a linear model analyzing the relationship between prediction error and key sweep parameters, $s$, which influences $t_f$, and sweep age $t_a$, which influences the degradation of the sweep signal.
